# Structure of the ILT Mutant Shaker K^+^ Channel and the Mechanism of Voltage Activation

**DOI:** 10.1101/2025.11.13.687653

**Authors:** Richa Agrawal, Ramon Mendoza Uriarte, Bernardo Pinto Anwandter, Trayder Thomas, Lydia Blachowicz, Young Hoon Koh, Francisco Bezanilla, Eduardo Perozo, Benoît Roux

## Abstract

To understand the process of voltage activation in Kv channels, we determined the cryo-EM structure of the Shaker channel ILT mutant (V369I, I372L, S376T), known to partially decouple the gating charge movement of the voltage sensing domain (VSD) and the opening of the pore domain (PD). The structure captures a previously unobserved intermediate state in which the VSD is only partially activated while the PD remains closed. Combined with computational modeling based on AlphaFold2 predictions and molecular dynamics simulations, it is shown that VSD activation does not mechanically pull the gate open, but instead induces a dynamic shift in the population equilibrium of the VSD–PD linker, providing the basis for electromechanical coupling between the two domains.

## INTRODUCTION

Voltage-gated potassium (Kv) channels are essential components of excitable membranes such as nerves and muscles, enabling the rapid and selective permeation of K⁺ ions in response to membrane depolarization (*2*). Kv channels are tetrameric membrane proteins in which each subunit contains 6 transmembrane helices (S1-S6) organized into two main structural features: the S1-S4 helices forming the voltage sensor domain (VSD) that responds to changes in the membrane potential, and the S5-S6 helices forming the pore domain (PD) that comprises the ion conducting selectivity filter and the intracellular activation gate (*3, 4*). In Kv channels, the VSD undergoes a series of conformational changes driven by the translocation of conserved arginine residues along the S4 helix, ultimately leading to pore opening (*5–8*). The conformational change within the VSD is thought to take place according to the sliding helix mechanism, which envisions that electrostatic forces hold down the S4 helix with its arginine gating charges at negative membrane potential, while membrane depolarization releases the S4 helix towards the extracellular side (*9–13*). However, the conformational changes underlying the electro-mechanical (E-M) coupling mechanism, by which the voltage-driven movement of the S4 helix in the VSD leads to the opening of the intracellular gate in the PD, are still being established. To resolve this issue, it is necessary to know the complete sequence of conformational states linking the closed resting state to the open activated state of a Kv channel.

The critical challenge to resolve the mechanism of voltage-activation in Kv channels is to experimentally capture all the relevant conformational states of the channel along the activation pathway. Efforts have been made to capture the structure of various Kv channels in different states (*14, 15*); however, the limited resolution did not provide sufficient information. Key information was obtained from structural studies of the Shaker K^+^ channel, a prototypical model of the Kv family. A cryo-EM structure was obtained of the Shaker WT channel in the open activated state, which is the dominant functional state at 0 mV (*16*). Recently, a cryo-EM structure of a mutant Shaker channel (I384R) that functionally uncouples the VSD from the pore domain was obtained, revealing a closed pore domain with a fully activated VSD (*17*). However, further information is required to elucidate the activation mechanism, as the molecular correlates linking VSD movement to pore opening remain unclear.

To resolve the mechanism of voltage-activation in Kv channels, the structure of the Shaker ILT triple mutant (V369I, I372L, S376T) in detergent micelles was determined using single particle Cryo-EM. These mutations localized along the S4 helix of the VSD alter the coupling between voltage-driven movements and gate opening. The latter remains closed even when moving ∼90% of the total gating charge with membrane depolarization, so that the channel is still closed at 0 mV and only a highly depolarized membrane potential succeeds to move the remainder of the gating charge and open the pore (**Fig. 1A**) (*18–20*). Despite functional studies characterizing the Shaker ILT mutant, as well as models of the mutations (*21, 22*), the structural basis for its functional phenotype remains unclear. For this reason, the ILT mutant provides a unique opportunity to capture a conformation of the channel near the final stages of voltage sensor movement and pore opening.

**Figure 1.**
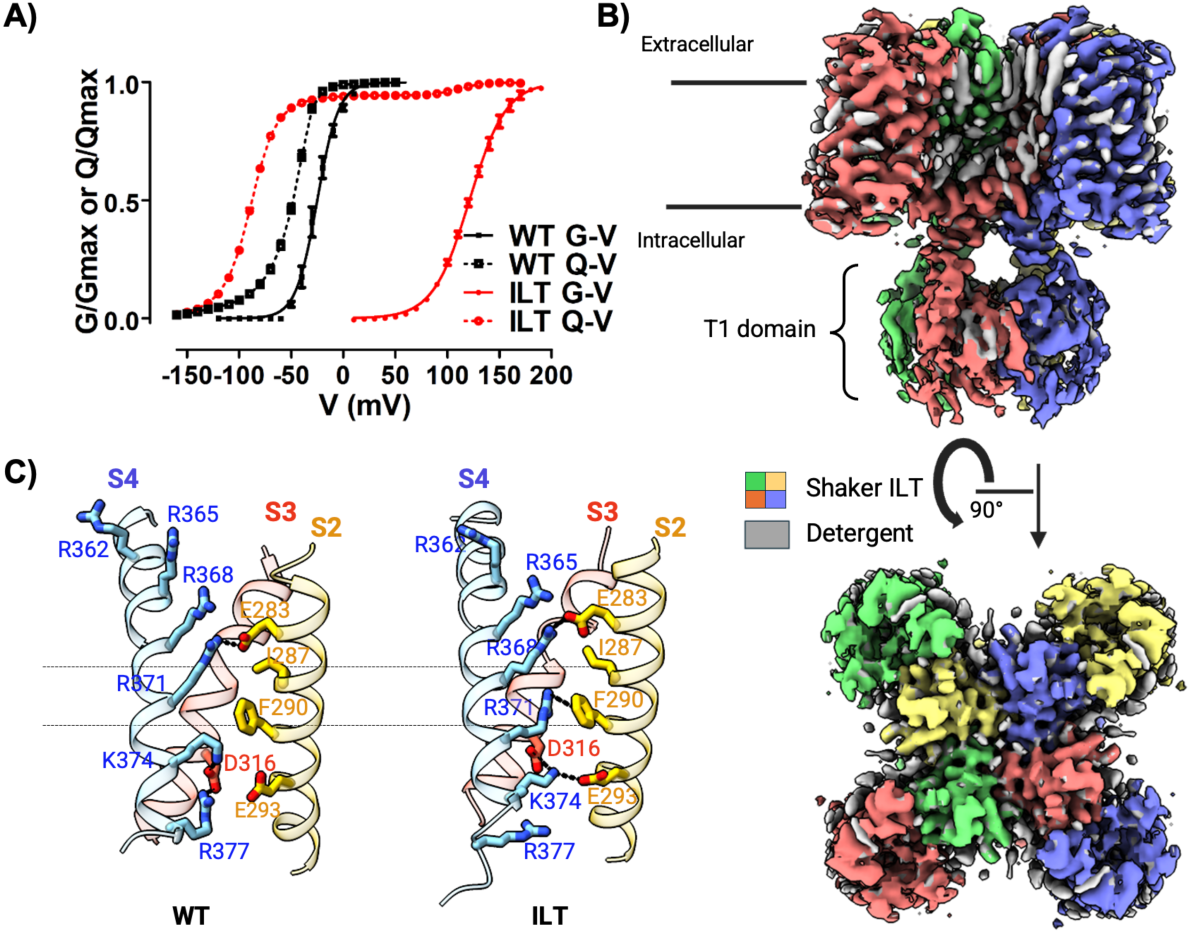
Structure of Shaker ILT. **A)** Normalized conductance (G/Gmax)-voltage curve (G-V) and normalized gating current Q/Qmax-voltage curve (Q-V); for Shaker WT (black) and Shaker ILT (red)**. B)** Side and top view of the tetramer map (sharpened) of Shaker ILT (PDB ID: 9ZS7). **C)** Gating charge interactions in the voltage sensor domain for both WT (left) and ILT (right). The S2 (gold), S3 (red), and S4 (cyan) helices are shown (S1 is not shown for the sake of clarity). The two dashed lines indicate the position of the hydrophobic region of the VSD., which is the main energy barrier in crossing the hydrophobic plug.

## RESULTS

### Structure of Shaker ILT

While the conductance vs. voltage relationship (G-V) of Shaker ILT mutant is strongly right-shifted to higher voltages compared to that of Shaker WT, most of the gating charge vs. voltage relationship (Q-V) is in fact left shifted (**Fig. 1A**) (*18, 19*). This was interpreted assuming that mutations stabilized the VSD in an intermediate state, with about 90% of the gating charge preceding the final transition to the conducting (open) state (*23*). The structure (**Fig. 1B** and additional information in **Figs. S1** and **S2**) was determined at a resolution of 3.4 Å, and it included density corresponding to the T1 domain, not previously reported. A hydrogen bond is present between the side chain of Arg487 near the end of the S6 helix and the main chain oxygen of Ile197 in the T1 domain of the same subunit (**Fig. S3**), suggesting a possible coupling between these structural elements. axis (**Fig. S4)**. Importantly, the inner bundle gate, ranging from V474 to N482, appears to be completely closed (**Fig. 2B**, **C and E**), in contrast to Shaker WT (**Fig. 2A** and **D**). The permeation pathway is constricted due to a rotation of the lower half of S6 with respect to the upper half of S6, centered around the second proline (Pro475) in the P-X-P hinge. There are some changes in van der Waals (VDW) interactions due to overall movement of the S5 and S6 helices towards the central axis of the pore. Labeling the four subunits A, B, C, D in the clockwise order from the extracellular side, residues Ala463(A) and Thr441(B) as well as Val467 and Thr442 (within the same chain) appear to make a stronger VDW contact in ILT compared to WT. The most significant constriction along the pore occurs at residues Ile470, Val474, Val478, and Asn482 (**Fig. 2**). This is confirmed by molecular dynamics (MD) simulation, as water molecules are prevented from entering into the pore (**Fig. S5**; **Fig. S6** shows a comparison of the pore radius among additional Shaker/Kv structures). Nonetheless, the selectivity filter of the Shaker ILT structure displays all the hallmarks of the conductive state, with no significant change compared to the Shaker WT structure (**Fig. 2F**). In particular, five K^+^ ions can be resolved in the binding sites (based on the sharpened map). The hydrogen bond interaction between Trp434 and Asp447 is intact (**Fig. S7**). In both Shaker WT and Shaker ILT, Trp435 of chain A forms a hydrogen bond with Tyr445 of chain B in the selectivity filter.

**Figure 2.**
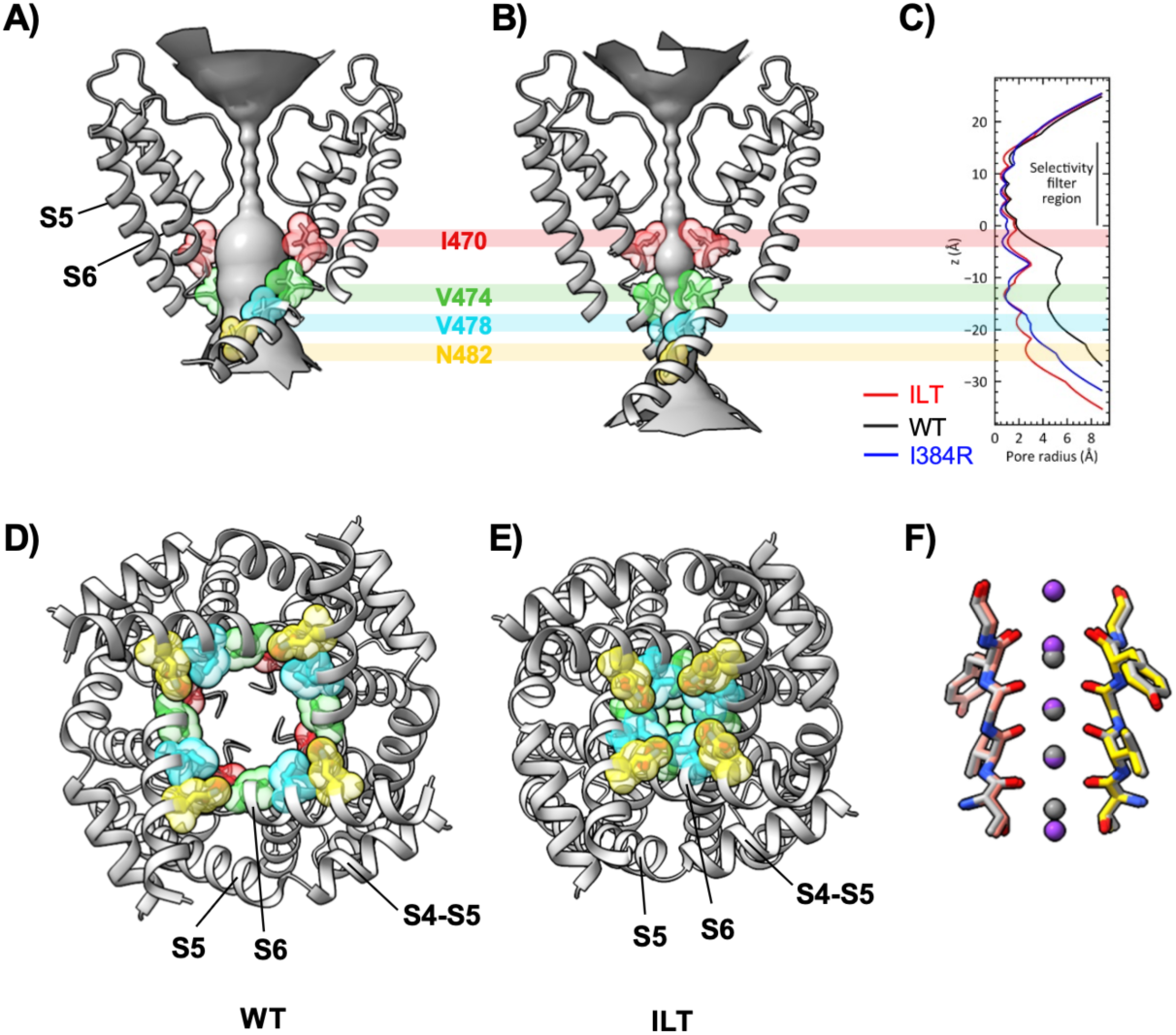
Analysis of the pore domain structure. A) and B) Side view of the WT (left) and ILT (right) pore domain. Four pore lining residues along the S6 helix responsible for constricting the permeation pathway in the ILT structure are highlighted in different colors. D) and E) Bottom view of the intracellular entrance to the permeation pathway showing the open gate for the WT channel and the closed gate for the ILT channel. The four pore lining residues are highlighted in the same colors as in panel A. C) Pore radius for the WT, I384R and ILT structures calculated from the HOLE program. (*1*) F) Overlay of selectivity filter region of WT (gray) and ILT (colored).

### Conformational changes in the voltage sensor domain (VSD)

The VSD in Shaker ILT displays important structural differences compared to WT (**Fig. 3A**). Consistent with the partial voltage activation observed in the Q-V curve at 0 mV (**Fig. 1A**), there is a downward shift of S4 by approximately one helical turn referred to as a click-down movement (*12, 24*). The downward shift of ∼5 Å is accompanied by a rotation of ∼15° (Fig. 3B) and an alteration of the pairing interactions of the gating arginine residues (**Fig. 1C**). In the WT structure, the outermost residue Arg365 is solvent-exposed, while the remaining gating charges form the interactions Arg368–Glu247(S1), Arg371–Glu283(S2), and Lys374–Asp316(S2). In contrast, in the ILT structure the gating charges form the interactions Arg365-Glu247(S1), Arg368-Glu283(S2), Arg371-Phe290(S2), and Lys374-Glu293(S2). More details about the change in interactions within the VSD are provided in **Table S1**. The VSD in the ILT structure is stabilized through a number of additional interactions with the PD (**Fig. 3C**). In particular, Phe373 in the S4 helix of subunit A forms a stacking interaction with Phe401 and Phe402 in the S5 helix of subunit C not observed in the WT structure (**Fig. S8**). The bottom of the ILT S4 helix is shifted towards the S5 helix of the adjacent subunit such that ILT-Phe373(S4) occupies a similar position to WT-Leu382 (S4-S5 linker) and now interacts with Phe401 and Phe402(S5). Of special importance are the interactions involving the three mutated residues (369, 372, and 376) with the adjacent S5 helix (**Fig. 3C**). In Shaker ILT, Ile369 interacts with Leu409 and Val408, while Leu372 interacts with Ile405. However, Thr376 does not appear to form robust interactions with adjacent residues. In Shaker WT, Val369 interacts with Leu409, while Ile372 interacts with Ile405, Val408, and Leu409. In addition, the backbone carbonyl oxygen of Ser376 forms a hydrogen bond interaction with the side chain of Gln383 in the S4-S5 linker.

**Figure 3.**
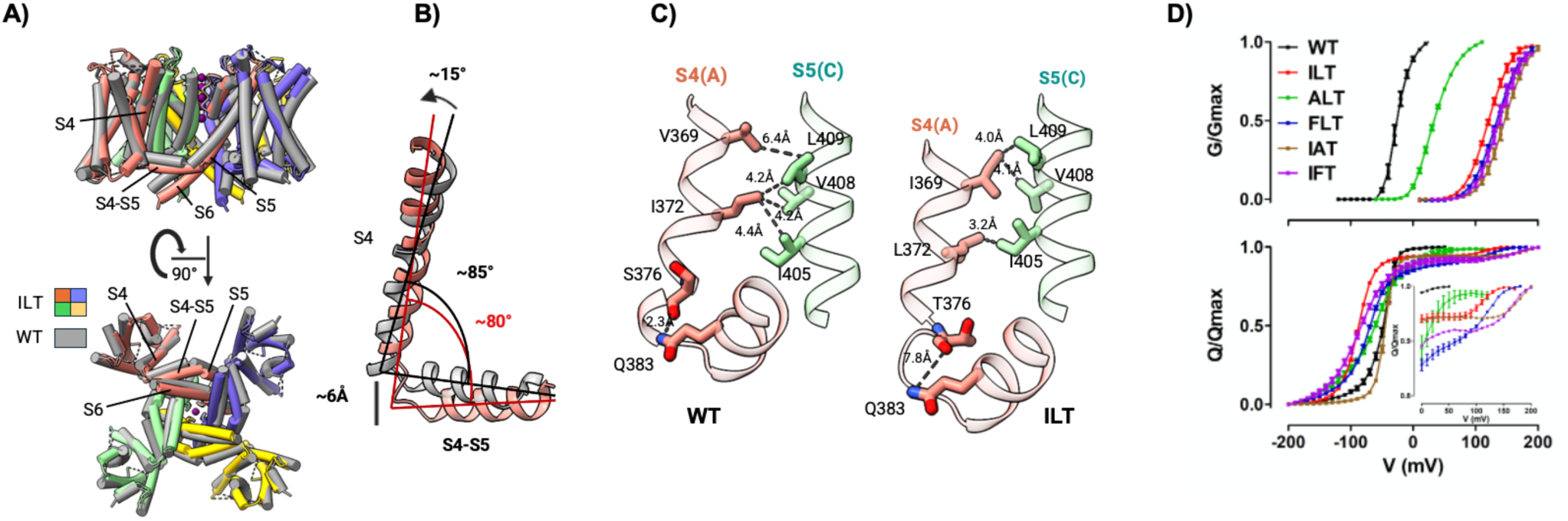
ILT Structure and interactions of ILT residues in the S4 helix: A) Front and bottom view of Shaker ILT structure (each chain shown in different color) superimposed with Shaker WT (gray); the T1 domain is not shown for the sake of clarity. **B)** The translational and rotational shift of the S4 helix and the downward shift of the S4-S5 linker in ILT (salmon) compared to WT (gray). **C)** ILT residues I369, L372 and T376 in the S4 helix (salmon) of chain A interact with hydrophobic residues (I405, V408, and L409) in the S5 helix (pale green) of chain C. D) Normalized conductance (G/Gmax)-voltage curve (G-V) and normalized gating current Q/Qmax-voltage curve for WT (black), ILT (red), ALT (green), FLT (blue), IAT (brown), and IFT (purple) (Inset is zoom view of final gating charge movement).

To further assess the importance of the various interactions, we experimentally investigated the effect of substituting a small (Ala) or bulky (Phe) residue at positions 369 and 372, yielding the series of Shaker mutants ALT, FLT, IAT, and IFT in substitution of ILT. These mutants were evaluated based on ionic and gating current measurements to derive G-V and Q-V curves. Gating currents were measured using a non-conducting mutant (W434F), which was also used to characterize Shaker WT and ILT. Comparing the G-V curves of mutants at position 369 to that of ILT, ALT becomes conductive at a lower voltage (-90 mV shift in G-V) while FLT shows a +10 mV shift (**Fig. 3D**). Q-V curves in both sets of mutants are shifted +10 mV. However, the diagnostic biphasic component observed in ILT is absent for ALT but increased in FLT. This observation suggests that the size of the residue in position 369 affects the stability of the ILT intermediate conformational state captured by the original ILT mutant. G-V curves for mutants at position 372 show a +25 mV and +15 mV shift when compared with ILT, IAT and IFT, respectively (**Fig. 3D**). Their Q-V curves show a +30 mV shift for IAT and a 0 mv shift for IFT, and both curves display the biphasic component. To better interpret these results, the Q-V curves were fitted with gating charge calculated from MD simulations based on Scheme S1 (in Supplementary Documents).

### Conformational changes in the S4-S5 linker and interactions with the inner bundle gate

In the WT structure, the VSD and the S4-S5 linker are in a fully activated (up) position consistent with the open gate formed by the S5 and S6 helices. In contrast, the linker is positioned to interact with the closed gate and keep it shut in the ILT structure (**Figs. S8** and **S9**). We will refer to this conformation as the “locked” linker. Remarkably, despite the structural and functional differences between the open or closed conformations of the WT and ILT structures, most of the main interactions involving the S4-S5 linker and the S5 and S6 helices forming the gate (Ile384 and Thr388 with Phe481, Phe484, and Tyr485) are conserved, though there are small shifts in side-chain configurations. Each S4-S5 linker interacts simultaneously with two S6 helices (**Figs. S4** and **S8**). All intra- and inter-subunit interactions are reproduced clockwise in the tetramer. Hydrophobic interactions across different subunits observed in ILT (Leu382 and Leu385 in the S4-S5 linker of subunit A with Phe401 and Phe402 in the S5 helix of subunit C) are not observed in WT (**Fig. S8A**), whereas there are hydrophobic interactions between Leu398 and Phe402 in the lower part of the S5 helix in subunit A with Phe481 of the S6 helix from subunit B in the WT structure (**Fig. S8D**). In the ILT structure, Asn482 and Tyr485 in the S6 helix from subunit B interact with Ser479 and Asn480 in the S6 helix of subunit A, while in the WT structure Tyr485 interact with Arg394 and Glu395 of the S4-S5 linker of subunit A (**Figs. S8B**, **S8C**). These interactions suggest that the ILT conformation is stabilized compared to WT by substituting both hydrophobic and hydrogen bond interactions. Regarding the configuration of the linker and the gate, it is of interest to compare the ILT and WT structures with the structure of the I384R Shaker mutant (PDB ID: 9OIC), which also displays the phenotype of VSD-PD decoupling in electrophysiology (*17*). The gate is closed in both the I384R and ILT structures, whereas the VSD is in its activated (up) position in both the I384R and WT structures (**Fig. S9**). The conformation of the S4-S5 linker and S6 helix in the I384R structure is between that of WT and ILT though closer to WT.

### AlphaFold modeling and MD simulations

To achieve a more complete perspective of the voltage activation process, the structure of Shaker ILT was used as a template to generate models of additional intermediate states with AlphaFold2 combined with MD simulations (**Fig. S10)**. The models are based on the knowledge that the VSD can adopt a few discrete conformations (*12, 24*). Those conformations associated with shift in helix turns of S4 can be conveniently identified based on the number of gating charges along the S4 helix crossing the nonpolar region formed by residues Phe290 and Ile287 (hydrophobic gasket) (*25, 26*), referred to as “clicks”. The VSD in the ILT intermediate structure corresponds to 1-click down from the fully activated VSD in the WT structure which is used as a reference state (0-click). Accordingly, this yields a total of five states (up, 1-, 2-, 3-, and 4-click down) and their associated incremental gating charge.

Several models generated by AlphaFold produced conformations with 2-clicks down relative to the WT structure (**Fig. S11**). However, no additional conformations of the VSD with S4 in a lower position could be generated. Starting with the AlphaFold predictions, two additional conformations were generated from this model using MD simulations with an applied membrane potential. A simulation at -400 mV produced a conformation with a VSD that is 3-clicks down relative to the WT structure, and a simulation at -750 mV produced a conformation with a VSD that is 4-clicks down relative to the WT structure. Combining these MD conformations with the two experimental structures (WT and ILT) and the AlphaFold model led to a total of five conformational states of the VSD. Counting the gating charges starting from the WT (up) state, the conformational transitions of the VSD can be described as 1-, 2-, 3-, or 4-clicks down, where the ILT conformation is 1-click down and the 4-clicks down model is said to be in the “down” state **(Fig. 4A)**.

**Figure 4.**
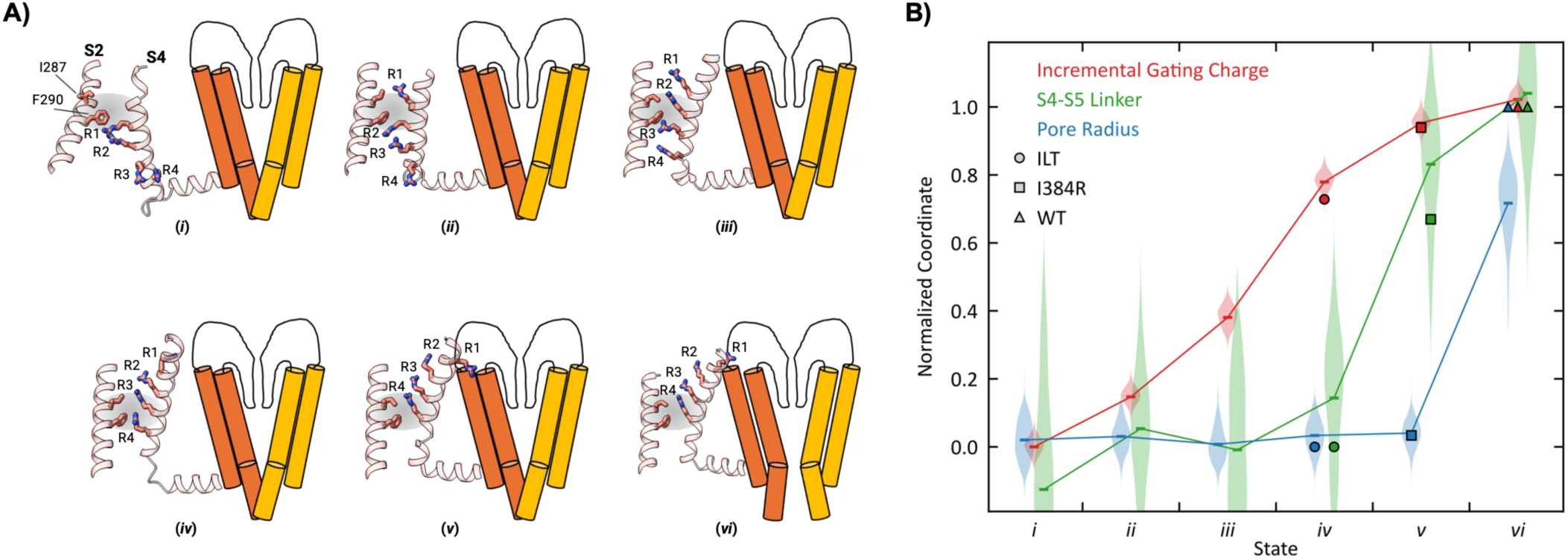
Conformational steps along the voltage activation of a Kv channel. **A)** The state of the Shaker channel is schematically depicted with 6 conformations (*i* to *vi*) in terms of 3 structural features: the gate of the PD, the VSD, and the S4-S5 linker. The gate can be either closed or open; the VSD can be in 5 discrete conformations; the S4-S5 linker can be locked (as in ILT) or activated (as in WT and I384R). State *iv* is the cryo-EM Shaker ILT structure (PDB ID: 9ZS7), state *v* is the I384R structure (PDB ID: 9OIC) and state *vi* is the WT structure (PDB ID: 7SIP). **B)** Relation between the incremental gating charge (red), the pore radius at Val474 (blue) and the position of the S4-S5 linker (green) for the 6 states along the activation pathway (coordinates have been normalized to fit between 0 and 1). The circle (ILT), square (I394R) and triangle (WT) represent the values from the three experimental cryo-EM structures. The violin plots represent the results from the MD simulations for each of the six states.

Special attention was dedicated to the conformational state between the ILT and WT structures, which is needed to elucidate the final stages preceding the opening of the gate after complete voltage activation. The three available cryo-EM Shaker structures (ILT, I384R, and WT) appear to map the final steps along the voltage activation pathway: ILT (VSD 1-click down, pore closed, linker locked) to I384R (VSD up, pore closed, linker activated), and then to WT (VSD up, pore open, linker activated). The cryo-EM structure of the Shaker I384R mutant, with its closed gate and fully activated VSD, is likely to be representative of this conformation. A first model of the intermediate state was generated from unbiased MD starting from the structure of the I384R mutant after reverting to the WT sequence. After 1 µs, all four S4-S5 linkers stabilized into the activated position that they occupy in the WT activated conformation while the gate remained closed. The final conformation is very close to the structure of Shaker I384R. A second model was generated through a targeted MD (TMD) simulation starting from Shaker-ILT (reverted to WT sequence), using the activated VSD S4 of Shaker WT as a target. TMD forces were applied only to the S4 helix, targeting the position of the fully activated (up) state of the VSD without restraint on the remaining residues (including the S4-S5 linker) followed by an unbiased MD simulation of 10 µs. During this process, two out of four S4-S5 linkers spontaneously made a transition from the locked position associated with the closed gate to the activated position that they occupy in the WT activated conformation (**Fig. S12** and **Movie S1)** while the two other linkers dissociate from the gate and become more dynamical, partly losing their helical structure. It is expected that, in time, all four S4-S5 linkers shall adopt the activated conformation while the intracellular gate remains closed. Therefore, in both cases, the state *v* corresponds to VSD(up)-PD(closed)-linker(activated) conformation.

In total, 6 potential conformational states of the channel are available (two experimental, four computationally predicted), labeled *i* to *vi*, mapping the activation process (**Movie S2).** Average features of the 6 different VSD conformation states are characterized from MD simulations in **Fig. S13**. In all these conformations, the inner bundle remains in a closed state. Interestingly, S4 adopts a standard α-helical conformation in the early states *i* to *iii* that progressively becomes a 3_10_ helical conformation in response to membrane depolarization beginning at state *iii* (**Fig. S13D**), in agreement with experiment (*27*). Analysis of MD trajectories of the 6 states representing the voltage activation process shows that Leu382 in the S4-S5 linker is more exposed to water molecules in the resting state but is inserted into a nonpolar pocket in the activated state, consistent with the experimental finding that substitution by a more polar (or less hydrophobic) residue stabilizes the resting state relative to the activated state, shifting the voltage-activation to higher voltages (*28, 29*). Furthermore, the gating charges of the individual activation steps of the channel calculated from MD simulations increase from 0 to 14.5 *e* (**Fig. 4B**) yielding families of Q-V curves that are in accord with electrophysiology measurements for WT as well as a series of mutants (**Fig. 3D** and **Fig. S14**). The fitted free energy of the 6 states shows the conformation observed in the cryo-EM structure of Shaker ILT (state *iv*) is stabilized relative to WT (state *vi*) by the triple mutations (**Fig. S14C**), hence explaining why it can be captured experimentally. This observation is consistent with the average change in interaction energy of the triple ILT mutation relative to WT calculated from the MD trajectories of the 6 states (**Fig. S10**). This analysis serves as validation of the structural models of the states along the voltage gating process.

## DISCUSSION

The voltage activation process of a Kv channel is schematically illustrated in **Fig. 4A** as a sequence of six conformational states (*i-vi*). The mechanism is inferred from an integrated analysis of three cryo-EM structures (Shaker ILT, I384R, and WT), electrophysiological measurements (**Fig. 3D**), structural modeling (**Figs. S10 and S11**), molecular dynamics simulations (**Figs. S12 and S13**), and kinetic modeling (**Fig. S14**). The structural elements playing a key role along the step-wise voltage activation pathway are the gate, the VSD and the S4-S5 linker. From state *i* to state *vi*, the gate goes from closed to open, the VSD progressively adopts five distinct conformations (0 to 4 clicks down from the WT up state), and the S4–S5 linker goes from a locked to an activated conformation. The incremental gating charge of each the six states along the voltage activation pathway yields a total of 14.4*e*, in good agreement with the experimentally observed value for WT Shaker (*7, 8, 30*). These incremental gating charges can be used to reproduce the measured Q-V of WT Shaker and several mutants based on a model assuming that the four VSD respond independently with a final cooperative step to the activated open state (**Fig. S14**), consistent with electrophysiology data (*30–35*) and the fact that each S4-S5 linker is simultaneously contacting two S6 segments (**Figs. S4** and **S8**).

Under a large negative potential, the resting state *i* is dominant and the gate is closed. **Fig. 4B** shows that voltage activation is an asynchronous process: the incremental gating charge increases progressively from state *i* to state *vi* (red line), but the S4-S5 linker only changes from the locked to the activated position in states *v* (green line) while the gate remains closed until it abruptly opens in the final transition to state *vi* (blue line). The changes through steps *ii* to *iv* (the conformation revealed by the cryo-EM structure of the Shaker ILT triple mutant) are strongly voltage-dependent with a total incremental charge of 13.7*e*. However, the S4-S5 linker remains in its locked position and the gate remains closed in states *ii*, *iii*, and *iv* despite the almost fully activated VSD. The movement of the last gating charge R371 along S4 across the hydrophobic plug is a requirement to open the gate (*25*), but even the transition from state *iv* to state *v*, in which the VSD becomes fully activated as in the I384R structure, does not readily cause the opening of the gate. In this state all four S4-S5 linkers adopt the activated (up) position seen in the WT structure, positioned to stabilize the open gate, which remains closed. In state *v*, the closed gate corresponds to a metastable conformation with a finite lifetime, ready to open spontaneously. The final transition from state *v* to state *vi* is only weakly voltage-dependent, with a small incremental charge of only ∼0.7*e,* presumably associated with the membrane electric field along the permeation pore—a scenario that is consistent with previous analysis of Shaker gating (*31–33*). The striking functional phenotype of Shaker ILT mutant is caused by the stabilization of the intermediate conformational state *iv* relative to Shaker WT (**Figs. S14** and **S15**). The stepwise voltage-activation process of Shaker ILT “stalls” in the intermediate state *v* even though the Q-V has almost reached its maximum while the G-V is still zero. Although the functional phenotype of Shaker ILT has traditionally been interpreted from the assumption that the mutations in the S4 helix decouple the movement of the VSD from the gate opening (*18–20*), the structural coupling between the VSD, the S4-S5 linker, and the intracellular gate is completely unaffected by the triple mutation.

In the canonical model of electromechanical coupling, VSD activation has been thought to mechanically pull the gate open. Rather, the final steps from state *iv* (ILT) to *v* (I384R) to *vi* (WT) show that the voltage-driven conformational change of the VSD progressively reshape the environment stabilizing the S4–S5 linker near the closed gate. In the penultimate pre-open state *v*, the linker shifts to its activated conformation, altering the open-closed equilibrium of the gate. Thus, the electromechanical coupling arises from a dynamic redistribution among metastable conformations of the linker rather than from direct mechanical leverage. We expect that this mechanism will be conserved in the channel superfamily.

## ACKNOWLEDGEMENT

The authors are grateful to the staff of the single particle Cryo-EM facility at The University of Chicago for their assistance during sample screening and data collection. The help of Patrick Haller is gratefully acknowledged. This work was supported by NIH grants 5R35GM152124 (BR), 5R01GM150272 (EP), R01GM030376 (FB), and by computational resources provided by the Research Computing Center of The University of Chicago and the Beagle3 high-performance GPU cluster funded by the NIH through grant 1S10OD028655.

**PDB deposited structure:** PDB ID: 9ZS7.

## MATERIALS AND METHODS

Detailed Materials and Methods can be seen in Supplementary Documents.

### Site directed mutagenesis and cloning

To generate the Shaker ILT construct, the mutations of V369I, I372L, and S376T were introduced by a modified QuikChange site-directed mutagenesis protocol (*36*) into the Shaker zH4 K+ channel with removed N-type inactivation (IR, Δ6-46) (*37*) in the pBSTA vector used for electrophysiological experiments. The mutation W434F was introduced on top of Shaker V369I/I372L/S376T (ILT) in pBSTA to record gating current (*38*). Other mutations (ALT, FLT, IAT, IFT) were generated for functional studies to measure gating and ionic current. For structural studies, Shaker ILT was cloned into the pEG BacMam vector containing an eGFP tag. All plasmid constructs used for expression were sequenced to confirm the desired mutations.

### Expression, purification, and structure determination

Shaker ILT was expressed in HEK293S GNTI-cells (ATCC) using the pEG BacMam plasmid and the Bac-to-Bac system (Invitrogen). Expressed protein was extracted from the membrane in the presence of 50mM Tris-Cl pH 7.5, 150mM KCl buffer, 2mM TECP (Buffer A) 1.5% DDM/0.15% CHS and purified in Buffer A containing 0.02% GDN. The concentrated protein (∼1 mg ml^−1)^ was immediately applied to UltrAufoil 300-mesh 1.2/1.3 Au grids (Quantifoil) and frozen in liquid-nitrogen-cooled liquid ethane. Grids were imaged at the University of Chicago Cryo-EM facility. Structure determination was performed using CryoSPARC (*39*)

### Electrophysiological recordings and analysis

pBSTA Shaker DNA was linearized using NotI enzyme and cRNA was transcribed using the T7 RNA kit (Invitrogen T7 Transcription Kit). cRNA was injected in defolliculated oocytes (stage V-VI) and incubated in SOS solution at 18 or 12 °C. After 1-4 days currents were recorded using the cut-open voltage-clamp method {Enrico, 1998 #131}. An in-house software (Analysis) was used to acquire and analyze the data. The individual K^+^ conductance were then normalized, averaged, and plotted against *V* to build conductance-voltage (G-V) curves. The data was fitted using two-state and three-state models (*23*).

### AlphaFold structure predictions

Structural models of the Shaker tetramer (residues 206-491) were generated using AlphaFold2 (*40, 41*)through ColabDesign v1.2.0b3 (*42, 43*). The coordinates of the S1/S5/S6 helices of the Shaker-ILT tetramer were used as template and initial guess. MSAs were generated for the VSD + S4-S5 linker and MSA reduction as per AF-Cluster (*44*). Structures were classified by VSD conformations and a representative structure with a 2 click-down VSD state was selected.

### MD Simulations

Simulations of the Shaker channel in the 6 conformational states were performed with OpenMM {Eastman, 2024 #74} using the CHARMM force field (*45–52*).The Shaker channel was inserted into a POPC bilayer and solvated with 150 mM KCl. After equilibration, production simulations were conducted for 1.2 µs. To explore the final stages of gate opening after voltage activation, TMD was performed to activate the VSD of the state *iv* structure. The resulting model with a closed gate and activated VSD was simulated unrestrained for 10 µs. To construct models of states *i* and *ii*, simulations were initialized from the equilibrated state *iii* system with an added electric field (*53*). Production runs were conducted with membrane potentials of -400 and -750 mV. Selected conformations with an individual VSD in 3 or 4 click-down states were assessed for stability without voltage, then the tetramer was symmetrized to the transitioned subunit by targeted MD (TMD).

### Gating charge calculation and Q-V modeling

The gating charge of Shaker in the 6 conformational states (WT and ILT) was calculated using MD simulations via the average displacement charge at zero membrane potential according to the Q-route theory (*53*). The experimental activation curve Q-V was fitted using the estimated gating charge transfer of the states (shown as the red curve in **Fig. 4B**) based on **Scheme S1**, which involves independent transitions of the four VSD through five states with the gate closed before a final cooperative transition to the open activated state (*31*). The free energies of each VSD state (relative to the activated state) were used as free parameters (**Fig. S14C**).

## MATERIALS AND METHODS

### Site directed mutagenesis and cloning

To generate the Shaker ILT construct, the mutations of V369I, I372L, and S376T were introduced by a modified QuikChange site-directed mutagenesis protocol (*1*) into the Shaker zH4 K+ channel with removed N-type inactivation (IR, Δ6-46) (*2*) in the pBSTA vector used for electrophysiological experiments. Additional mutations W434F were introduced on top of Shaker V369I/I372L/S376T (ILT) in pBSTA to record gating current (*3*). Other mutations (ALT, FLT, IAT, IFT) were generated for functional studies to measure gating and ionic current. For structural studies, Shaker ILT was cloned into the 5’ EcoRI and 3’ XbaI sites of a modified pEG BacMam vector containing a C-terminal HRV 3C protease site, an eGFP tag and an 8× His tag. All plasmid constructs used for expression were checked by either Sanger (University of Chicago DNA Sequencing Core Facility) or nanopore (Plasmidsaurus) sequencing to confirm the desired mutations.

### Cell lines

The HEK293S GnTI- cells in suspension that were used for protein expression and purification were obtained from ATCC (CRL-3022). GnTI- cells were grown at 37 °C and 7.8% CO_2_ in FreeStyle 293 expression medium (Gibco, Thermo Fisher Scientific) supplemented with 2% heat-inactivated fetal bovine serum (FBS) and 10 µg ml^−1^ penicillin–streptomycin. Sf9 cells (Thermo Fisher Scientific, 12659017) were cultured in SF-900 II SFM medium (Gibco, Thermo Fisher Scientific) supplemented with 10% FBS and 10 µg ml−1 gentamicin at 28 °C.

### Expression and purification of Shaker ILT

To obtain a biochemically stable preparation suitable for single-particle cryo-electron microscopy (Cryo-EM), the Shaker ILT protein was expressed in HEK293S GNTI- cells (ATCC). The pEG BacMam/Shaker ILT plasmid and the Bac-to-Bac system (Invitrogen) were used to generate a recombinant bacmid. Sf9 insect cells (Thermo Fisher Scientific) were transfected with the bacmid using Cellfectin II (Thermo Fisher Scientific) to generate P0 baculovirus. After amplification, the P1 baculovirus was used to transduce HEK293S GNTI- cells at a ratio of 1:10 (v/v). After 20–24 h of incubation at 37 °C, 10 mM sodium butyrate was added to the cells and the culture was transferred to 30 °C. Cells were collected 54–58 h after transduction by low-speed centrifugation. The cell pellets were washed with phosphate-buffered saline pH 7.4, collected by low-speed centrifugation, flash-frozen and stored at −80 °C for later purification. Purification was carried out on ice or at 4 °C. The flash-frozen cell pellets (0.5 l) were thawed in a water bath at room temperature and Dounce homogenized in the presence of 50mM Tris-Cl pH 7.5, 150mM KCl buffer, 2mM TECP (Buffer A) containing 10 µg ml−1 DNase and a complete Protease Inhibitor Cocktail (Roche). Protein was extracted with a 1.5% n-dodecyl-β-d-maltopyranoside (DDM; Anatrace) and 0.15% cholesteryl hemisuccinate (Anatrace) for 120 min. Solubilized supernatant was isolated by ultracentrifugation at 100,000xg RPM, 1h, 4°C. The supernatant was incubated for 2 h with 2 ml CNBR-activated Sepharose beads (GE Healthcare) coupled with 4 mg high-affinity GFP nanobodies. The beads were transferred to a plastic column and further washed with 10 column volumes of buffer A containing 0.05% DDM (Anatrace) and 0.01% cholesteryl hemisuccinate, exchanging stepwise to buffer A containing 0.02% GDN. Protein was cleaved by HRV 3C protease overnight at 4 °C, concentrated and analysed using SEC on a Superose 6, 10/300 GE column (GE Healthcare) in buffer A with 0.02% GDN. Peak fractions were collected and concentrated using a 100 kDa molecular mass cut-off centrifugal filter (Millipore) to ∼1 mg ml^−1^. The concentrated protein was used immediately for cryo-EM.

### Cryo-EM sample preparation and data acquisition

UltrAufoil 300-mesh 1.2/1.3 Au grids (Quantifoil) were plasma-cleaned at 20W for 40 s in an air mixture in a Solarus Plasma Cleaner (Gatan). Purified Shaker ILT samples were applied to the grids and frozen in liquid-nitrogen-cooled liquid ethane using a Vitrobot Mark IV (FEI) and the following parameters: 3.5 µl sample volume; 3.5 s blot time; 1.0 blot force; 100% humidity; temperature of RT. Grids were screened for single particles on a 200kV Glacios TEM with cryo-autoloader equipped with a K2 summit direct detector (Gatan). Grids were imaged at the University of Chicago on a Titan Krios with a K3 detector and GIF energy filter (set to 20 eV) at a nominal magnification of ×81,000, corresponding to physical pixel size of 1.068 Å. Movies were acquired for 50 frames with total exposure 60 e^−^ A^−2^.

### Single-particle cryo-EM analysis

All steps for structure determination were performed using CryoSPARC (*4*), including motion correction and contrast transfer function (CTF) estimation. A dataset of ∼6600 movies was collected, and particles were picked and classified in 2D to generate templates for template-based particle picking. Approximately 2.6 million initial particles were picked and subjected to 2 rounds of 2D classification. From these, 0.45 million particles were selected to generate two *ab initio* models for C1 and C4 symmetry separately. This process was repeated multiple times to eliminate junk particles. ∼0.29 million particles from the best class were processed for 3D refinement (non-uniform or local refinement) with the C4 refinement algorithm (*5*), which consistently yields the best results. Reference-Based Motion correction was applied to estimate per-particle movement trajectories and empirical dose weights. These particles underwent a further 3D refinement step, followed by local refinement using a mask with enforced C4 symmetry. Local resolution was calculated using CryoSPARC.

### Model building, refinement and molecular visualization

The Shaker-IR WT structure (PDB ID: 7SIP) was used as a template to build atomic models for Shaker ILT. Secondary structural elements were fit into the map and further model building was pursued through iterative rounds of manual model building in COOT (*6*) registering secondary structural elements using bulky residues such as Phe and Arg. Simultaneously, flexible loops fitting in position in the atomic models were obtained using interactive flexible fitting in ISOLDE (*7*). The density was of sufficient quality to assign rotamers for key residues whereas side chains of some residues that could not be assigned even tentative rotamers were truncated at the Cβ position of the residue. The tetramer model was generated by applying C4 symmetry operations to the monomer in UCSF ChimeraX (*8, 9*). Models were refined iteratively using phenix.real_space_refine in Phenix (*10*). PDB validation has been performed in the PDB validation server. All structural analyses and figure generation were performed using UCSF ChimeraX.

### Electrophysiological recordings and analysis

*Xenopus laevis* ovaries were obtained from Xenopus 1 (Dexter, Michigan). The follicular membrane was digested by collagenase 2 mg/mL supplemented with bovine serum albumin 1mg/mL. Oocytes were kept at 12 or 18 °C in SOS solution containing (in mM) 96 NaCl, 2 KCl, 1 MgCl_2_, 1.8 CaCl_2_, 10 HEPES, pH 7.4 (NaOH) supplemented with gentamicin (50 mg/ml). We used clones of the Shaker zH4 K^+^ channel with removed N-type inactivation (IR, Δ6-46) in the pBSTA vector (*2*). pBSTA Shaker DNA was linearized using NotI enzyme and cRNA was transcribed using the T7 RNA kit (Invitrogen T7 Transcription Kit). cRNA was injected in defolliculated oocytes (stage V-VI) and incubated in SOS solution at 18 or 12 °C. After 1-4 days currents were recorded using the cut-open voltage-clamp method (*11*). Voltage-sensing pipettes were pulled using a horizontal puller (P-87 Model, Sutter Instruments, Novato, CA). The resistance ranged between 0.2-0.5 MΩ. Data were filtered online at 20–50 kHz using a built-in low-pass four-pole Bessel filter in the voltage clamp amplifier (CA-1B, Dagan Corporation, Minneapolis, MN, USA) sampled at 1 MHz, digitized at 16-bits and digitally filtered at Nyquist frequency (USB-16042AO; Measurement Computing, Norton, MA) using Gpatch64M (in-house software). An in-house software (Analysis) was used to acquire and analyze the data. External solution for ionic recording was composed of (mM): K-Methanosulfonate (MES) 12, N-Methyl D-glucamine (NMG)-MES 108, Ca-MES 2, HEPES 10, pH 7.4, and internal solution of (mM): K-MES 120, EGTA 2mM, HEPES 10, pH7.4. For right shifted mutants the external solution contained K-MES 120, Ca-MES 2, HEPES 10. External solution for gating currents recording was composed of (mM): NMG-MES 120, Ba-MES 2, HEPES 10, pH 7.4, and internal solution of (mM): NMG-MES 120, EGTA 2mM, HEPES 10, pH 7. The ionic current was measured when the current reached a steady-state level and then converted to conductance (G) using the equation, *G* = *I*⁄[*V* − (*RT*⁄*F*) ln([*K*^+^]^out^⁄[*K*^+^]_in_)], where *I* is the current, *V* is the membrane voltage, *R* is the gas constant, *T* is the temperature in Kelvin, *F* is the Faraday constant, and [K^+^]_in_ and [K^+^]_out_ are the intracellular and extracellular K^+^ concentrations, respectively. The individual K^+^ conductances were then normalized, averaged, and plotted against *V* to build conductance-voltage (G-V) curves. The data was fitted using two-state and three-state models (*12*).

### AlphaFold structure predictions

Structural models of the Shaker tetramer (residues 206-491) were generated using AlphaFold2 (*13, 14*) through ColabDesign v1.2.0b3 (*15, 16*). The coordinates of the S1/S5/S6 helices of the Shaker-ILT tetramer in the intermediate gate-closed conformation were used as the only structural template and initial guess. MSAs were generated for the VSD + S4-S5 linker (residue 206-397) in either WT or ILT sequences using the ColabFold 1.5.5 API and the AF-Cluster (*17*). MSA reduction method was used to cluster the MSA (39 clusters for WT, 32 clusters for ILT) to prompt exploration of alternative VSD conformations. Each MSA sequence was then recombined with the pore-domain sequence into a full-length sequence. Each MSA cluster and the pore-domain template was then input into AlphaFold, generating a structure for each cluster and each of the 5 AlphaFold2 multimer_v3 models. Each structure was recycled a maximum of 20 times, using the coordinates of the previous cycle, an early stop tolerance of 0.5 Å RMSD between cycles, and with no activation of dropout layers. Misfolded structures were filtered out and the remaining 312 out of 355 structures were then classified into 0, 1, or 2 click-down states by whether the sidechain of each S4-helix Arg residues was below the hydrophobic plug formed by residue F290 and I287. Of 7 structures categorized as 2-click down, 1 had an S4 shifted down by a full turn and was selected for MD simulations as described below.

### MD simulations

Atomic models for the Shaker channel in various conformational states were constructed using residues 206 to 491 for MD simulations. The simulations were performed with OpenMM 8.1.2 (*18*). The CHARMM force field CHARMM36m(*19*) for protein (*20–23*), lipids(*24*), ions(*25*), and TIP3P(*26*) was used. The default parameters from the CHARMM-GUI input files were used except a switching function was used to smoothly truncate the potential at the cutoff beginning at 10 Å. The temperature was held constant at 310 K. Constant pressure simulations were performed at 1 atm. An initial system containing the Shaker channel was first prepared using the CHARMM-GUI membrane builder,(*27–30*) which is inserted in a bilayer of POPC (1-palmitoyl-2-oleoyl-phosphatidylcholine) lipids and solvated in 150 mM KCl, and with all sidechains being assigned their standard protonation state. Subsequently, the channel can then be swapped with its other conformational states. In all cases the selectivity filter was initially loaded with K^+^ ions at binding sites S_0_, S_2_, and S_4_,(*31*) with water occupying the intervening sites.

Simulations were conducted at a 2 fs timestep and with a force constant of 1 kcal/mol/Å^2^ for positional restraints, except when stated otherwise. Equilibration was generally performed in 3 steps, with changes noted below. In all steps the loops of the VSD (at the level of Thr265 and Ala343 Cα atoms) were held packed in a 50 Å radius cylinder centered on the channel to prevent the full extension of the loops in the xy-plane. The 1st equilibration step was performed with positional restraints on all protein residues, excluding those that are unresolved in the cryo-EM structure, and a flat-bottom restraint applied to the oxygen of water molecules to prevent them from entering the membrane region. The system was energy minimized for 400 steps and then equilibrated for 100 ps at NVT (constant Number of particles, Volume, and Temperature) conditions, followed by 10 ns at NPT (constant Number of particles, Pressure, and Temperature) conditions. A 2nd equilibration step was performed without the water restraints for 20 ns or until the box-volume converged, taking at least 90 ns but no more than 130 ns. After this step, the channel is reverted/mutated to a common sequence, in either WT or ILT, and the gating charge is determined as detailed below. Next, the 3rd equilibration step was performed for 10 ns with protein restraints reduced to backbone restraints on all TM helices, S4-S5 linker, pore helix, and selectivity filter. Production simulations were then performed for 1.2 µs with a center of mass (COM) positional restraint on Val467 Cα atoms to prevent the channel from drifting. The trajectories were analyzed with VMD (*32*) and the first 200 ns were omitted from the analysis.

For the simulations based on the cryo-EM structures, the protein was restrained during the 1st equilibration step with a force constant of 100 kcal/mol/Å^2^. The 3rd step was performed for 11 ns with the 100 kcal/mol/Å^2^ position restraint being linearly scaled off over the last 10 ns. The state *ii* conformation in WT sequence was only equilibrated following the 1st step, and with only the Cα atoms restrained.

A model of Shaker WT with a closed gate (as in ILT) and activated VSD (as in WT) was generated to explore the final stages of gate opening after voltage activation using targeted MD. The TMD is started from Shaker ILT reverted to WT after the 2nd equilibration step and the activated VSD S4 of 7SIP is used as a target. After reverting the mutant, the system was first energy minimized for 200 steps, and equilibrated for 50 ns at NPT conditions with the backbone restrained. The activated VSD S4 was then targeted starting at a 1 kcal/mol/Å^2^ restraint that was linearly increased to 100 kcal/mol/Å^2^ over 11 ns while the remaining residues (including the S4-S5 linker) were left free. This was then equilibrated for 10 ns with only the S4 helix backbone restrained at 1 kcal/mol/Å^2^. A production simulation was then taken to 10 µs with the S4 released.

### Transmembrane potential simulations

A model for the resting state of Shaker, closed gate and VSD down, was explored with a transmembrane potential at hyperpolarizing voltages, which was implemented with an electric field (*33*). The Shaker ILT in ILT and WT sequence and the Shaker WT AlphaFold2 model were each considered at -400 and -750 mV. All systems were initialized after the equilibrations described above plus an added 100 ns equilibration at NPT conditions with backbone restraints on the PD, secondary structure restraints on the S4-S5 linker, and the initial loading state of the selectivity filter held fixed with position restraints on the K^+^ ions. The VSD was left free to move. Afterwards the membrane potential at - 400 or -750 mV was introduced at NVT conditions. The first 200 ns were done at 2 fs, which was then increased to 4 fs after hydrogen mass repartitioning, the hydrogen mass was increased threefold. The simulations at -400 mV were sampled for 6 µs and simulations at -750 mV for 3 µs. A subunit was observed to transition to state *ii* for the ILT sequence after 1.6 µs at -750 mV, and for the AlphaFold2 model after 400 ns at -400 mV and to state *i* after 2 µs at -750 mV. To assess the reliability of the obtained conformation of state *i* from the AlphaFold2 model at -750 mV an initial equilibration was performed at NPT conditions without voltage for 200 ns. Next, a TMD step is used to obtain the final model because all VSDs act independently of the others so not all subunits complete the transition; thus, in the TMD the VSD and S4-S5 linker residues of the subunit that completed the transition is used as a target to the remaining subunits. The PD was restrained in the closed gate conformation in this TMD step. The TMD restraint began at 1 kcal/mol/Å^2^ and is linearly increased to 100 kcal/mol/Å^2^ over 11 ns at NPT conditions. The channel was then symmetrized in vacuum for 55 ns at NVE (constant Number of particles, Volume, and Energy) conditions with the Verlet integrator at a 1 fs timestep. The symmetry restraint began at 1 kcal/mol/Å^2^ and was linearly increased to 100 kcal/mol/Å^2^ in the first 11 ns. The final model was acquired after 2,000 steps of minimization with the symmetry restraint increased by four-fold. The final models were then assessed with production simulation as described above.

### Gating charge calculation

The gating charge of Shaker in the six conformational states was calculated using MD simulations via the average displacement charge at zero membrane potential according to the Q-route theory (*33, 34*). After the 2nd equilibration step, each state was then mutated to a common sequence (WT or ILT). The channels requiring a mutation were first energy minimized for 200 steps. The simulations to calculate the gating charge were performed at NPT conditions with backbone restraints on all resolved residues and the initial occupancy of the selectivity filter held fixed with restraints on the K^+^ ions z-position. Three replicates were performed for each state; WT sequences were simulated for 50 ns and ILT sequences for 20 ns. The gating charge was then estimated from the displacement charge, omitting the first 10 ns from the calculation (*33, 34*).The experimental activation curve Q-V was fitted using the estimated gating charge transfer of the states. The activation process is represented according to **Scheme S1**,

**Scheme S1.**
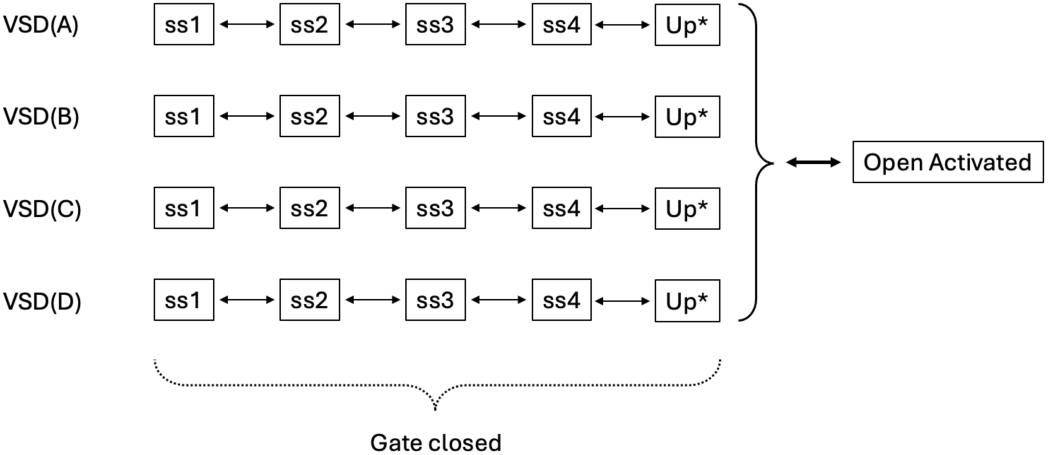

which involves independent transitions of the four VSD through five states with the gate closed before a final cooperative transition to the open activated state. The equation for the total gating charge *Q* (expressed in elementary charge) as a function of voltage *V* follows,

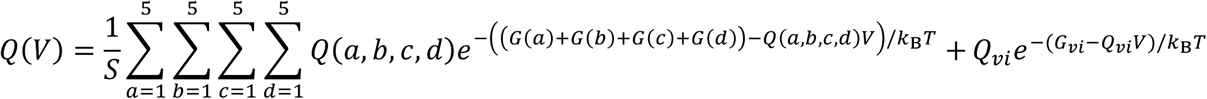

where *Q_vi_* is the incremental charge of the state *vi* (open activated), and the index *a, b, c, d* denote the states of each of the four VSD, yielding the incremental charge

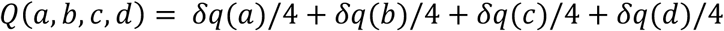

with the normalization constant,

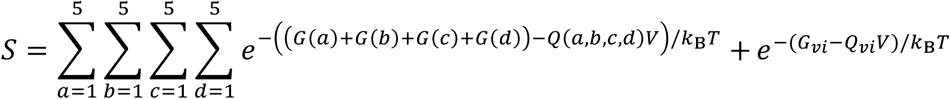

The Q-V curves are normalized to 1 at large voltage. This scheme is reminiscent of a combination of class B and D kinetic models proposed in Zagotta et al.(*35*), which states that each subunit must undergo four conformational changes at the VSD before opening with a final concerted opening of the gate (*36*). The free energy of each closed state was fitted relative to the free energy of the activated state (fixed to zero). The gating charge estimates from the MD simulations and their uncertainty were used in the fitting. The results are shown in **Fig. S13**.

**Figure S1.**
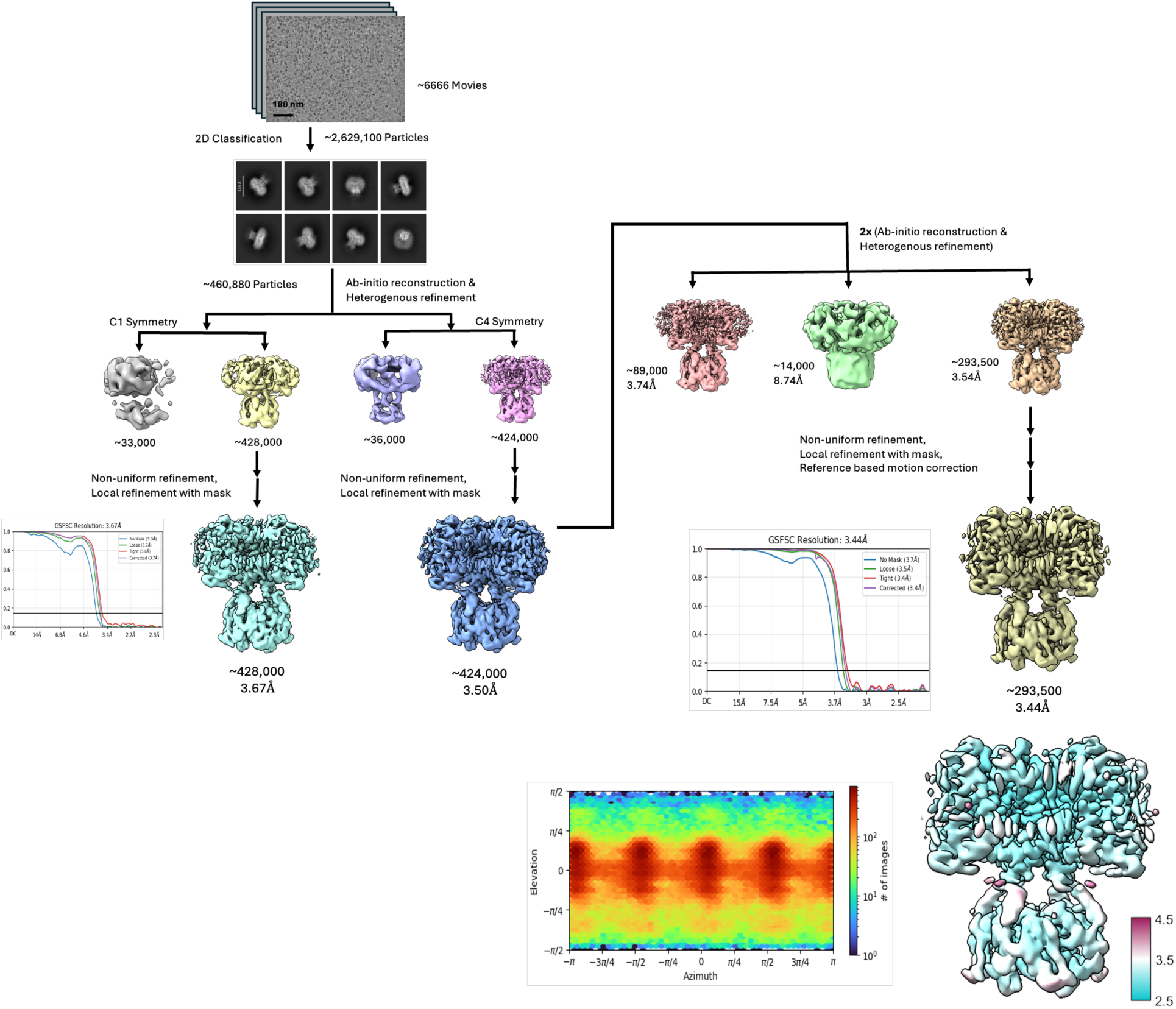
Data processing of Shaker ILT. Data processing was done using CryoSPARC for motion correction, 2D classification, 3D classification and non-uniform refinement. Final resolution of map estimated to be 3.4 Å and local resolution for transmembrane region estimated to be ∼3.0 Å.

**Figure S2.**
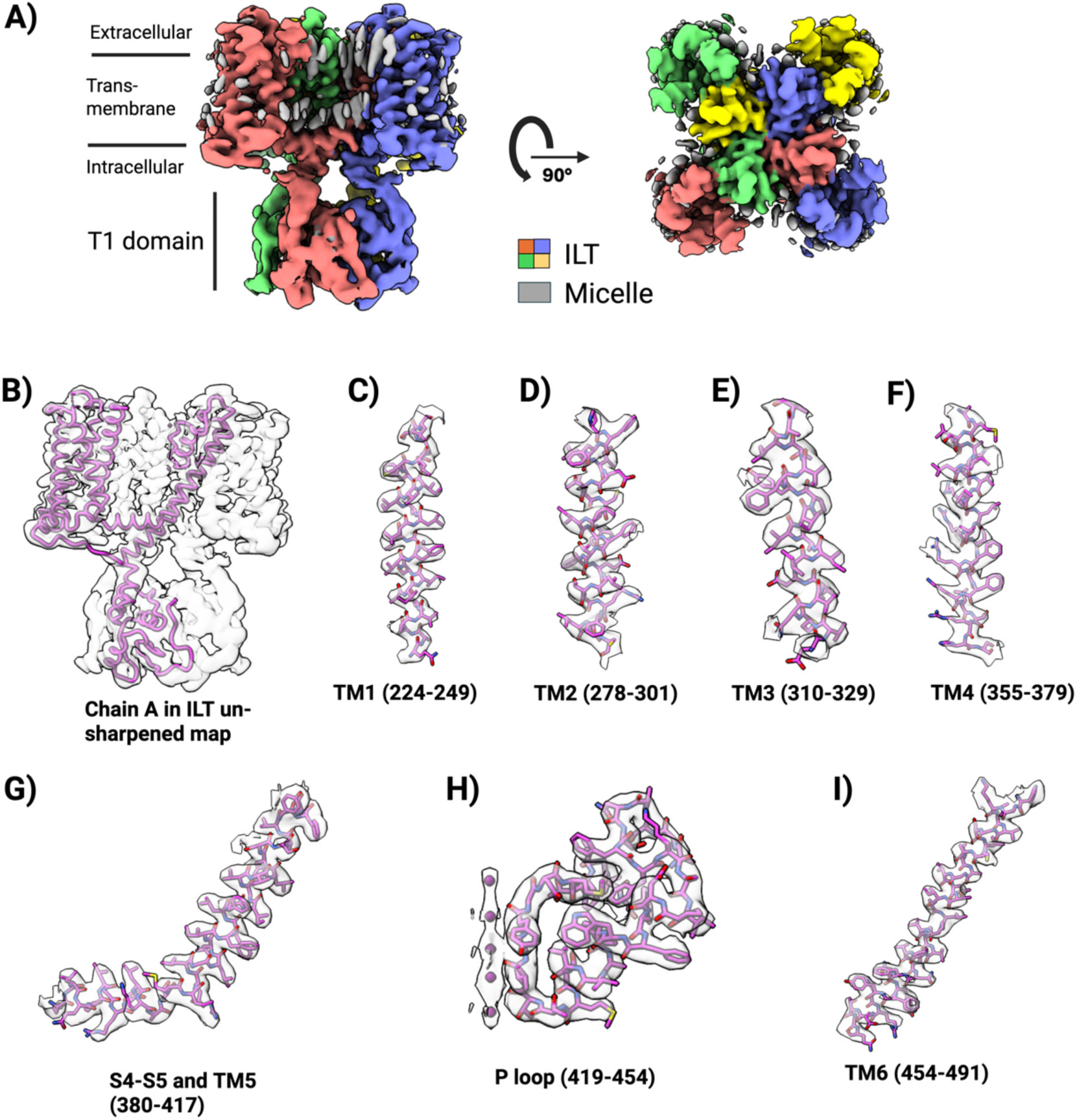
Map density of ILT structure. **A)** Front and top view of unsharpened map of Shaker ILT with each chain shown in a different color. **B)** Fitting of monomer chain in Shaker ILT map density. **C-F)** Transmembrane helices from S1-S4 of voltage sensor domain with atomic coordinates. **G&I)** S4-S5 linker, S5 and S6 helices of pore domain with atomic coordinates. **H)** Pore helix and selectivity filter with atomic coordinates including potassium ion density (based on sharpened map).

**Figure S3.**
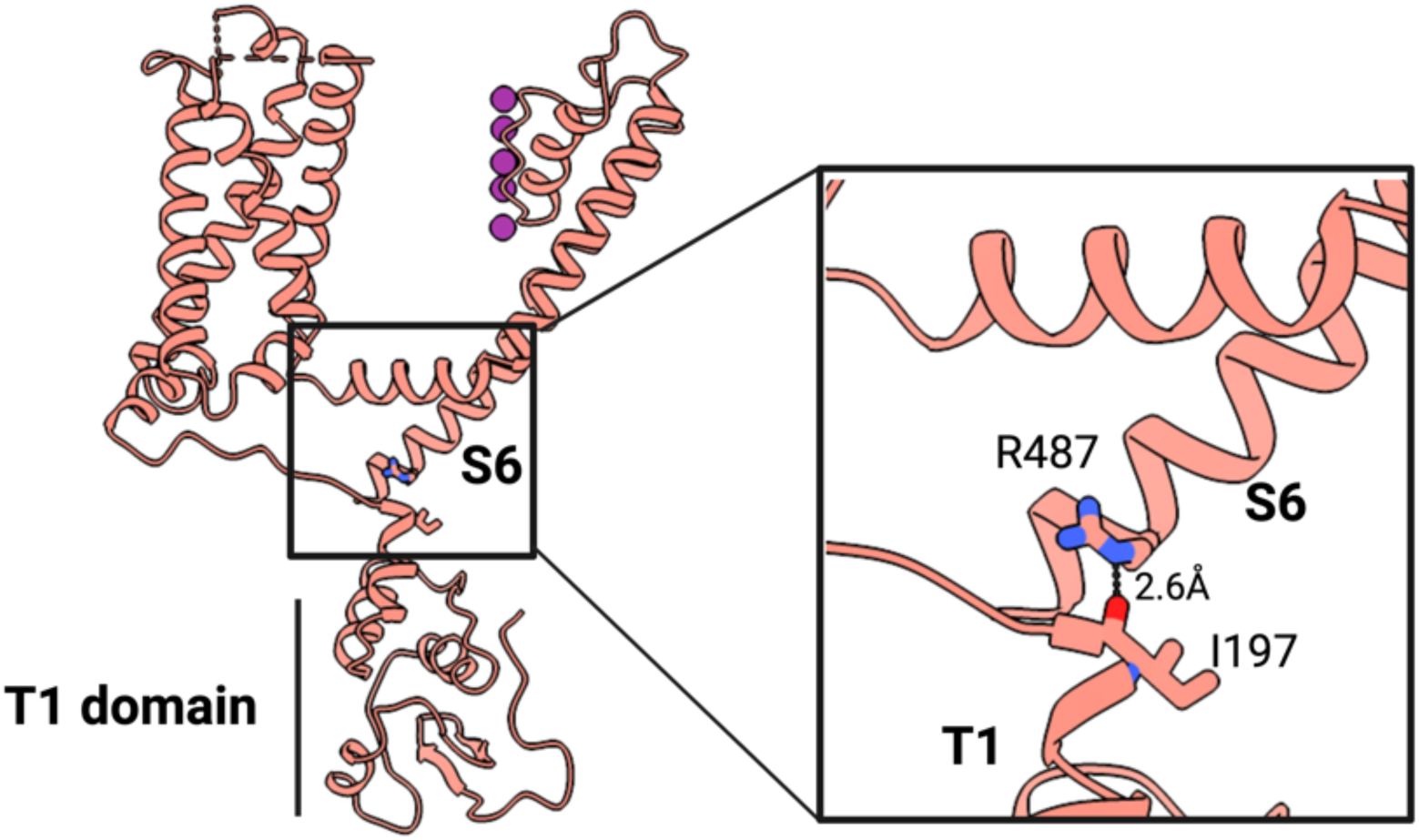
T1 domain interaction with S6 helix. Chain A of tetramer showing carbonyl oxygen of I197 in T1 domain interacting with R487 in S6 helix of pore domain; zoomed view in the inset.

**Figure S4.**
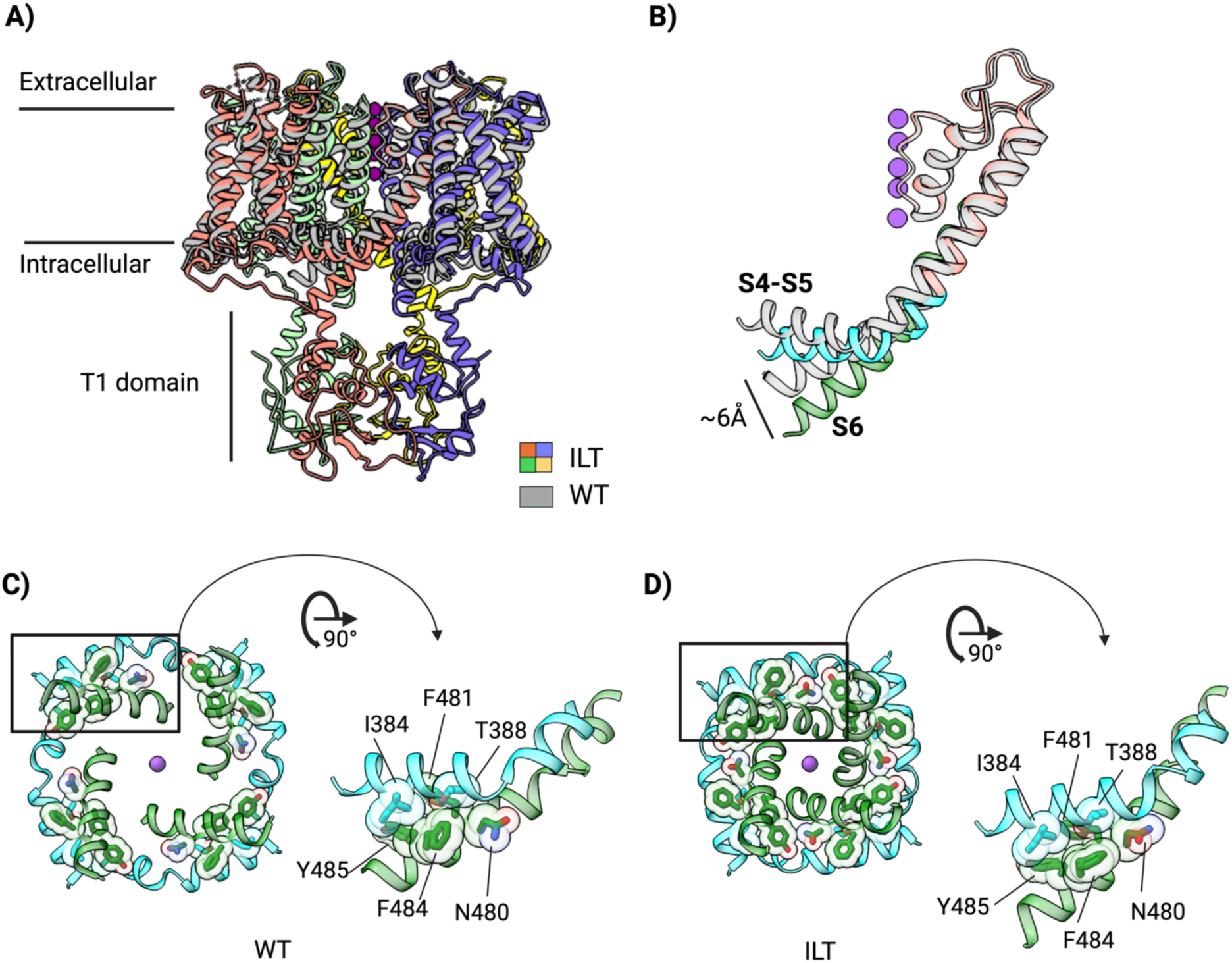
Movement of S6 helix relative to S4-S5 linker. **A)** Ribbon representation of Shaker ILT and WT tetramer. **B)** Shift in S4-S5 linker (cyan) and S6 helix (green) of ILT compared to WT (gray). **C)** Bottom and front view showing interactions of S4-S5 linker (cyan) and S6 helix (green) in WT. **E)** Bottom and front view showing interactions of S4-S5 linker (cyan) and S6 helix (green) in ILT.

**Figure S5.**
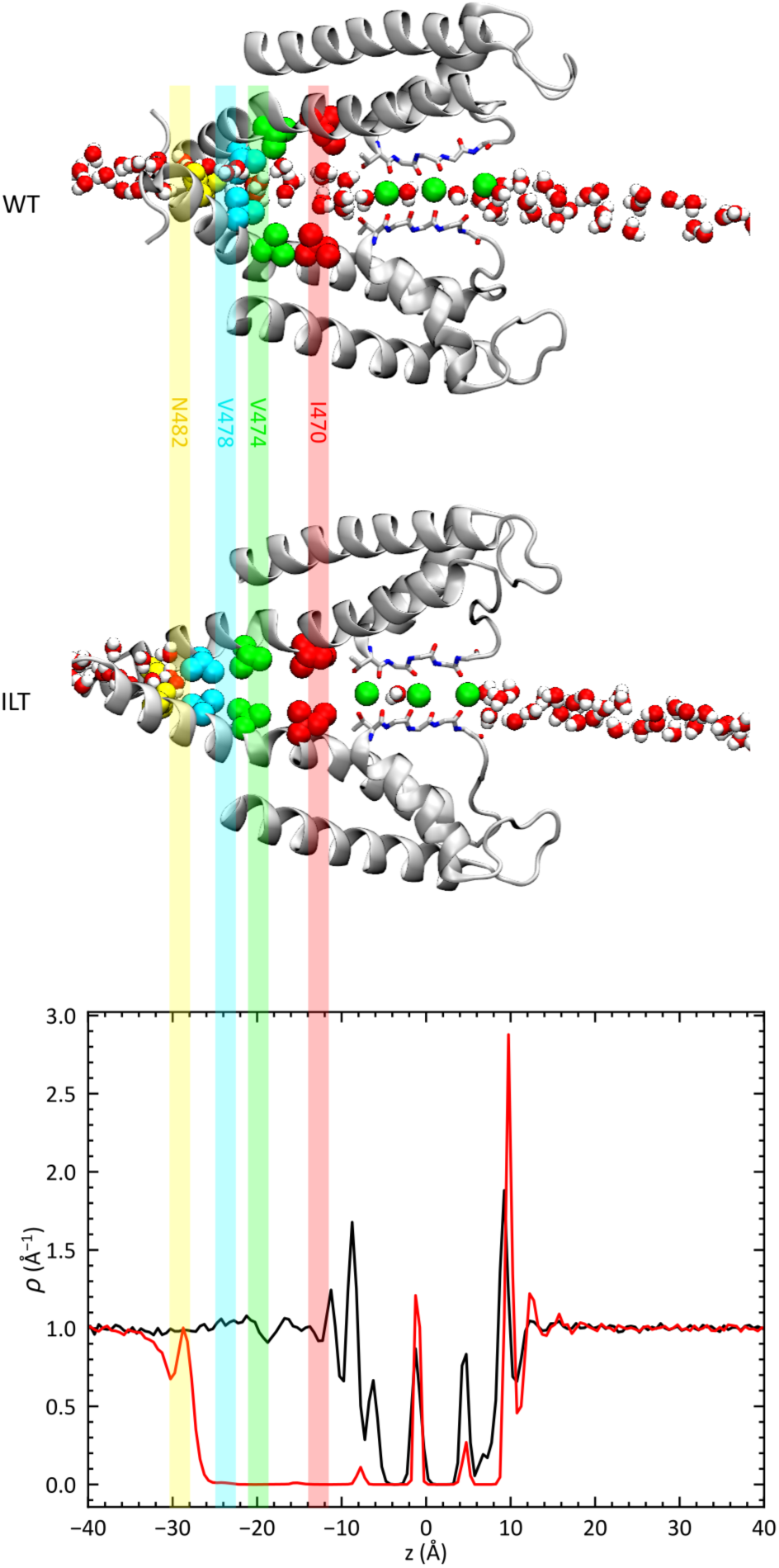
Water profile along the pore axis of Shaker. At the top are shown the final snapshots from the 1.2 µs trajectories for WT (PDB ID: 7SIP) and ILT (PDB ID: 9ZS7) showing the water within 2.5 Å from the pore axis. At the bottom are shown the line densities of water within 2.5 Å of the pore axis relative to the bulk along z-axis from molecular dynamics simulation in WT (black) and ILT (red).

**Figure S6.**
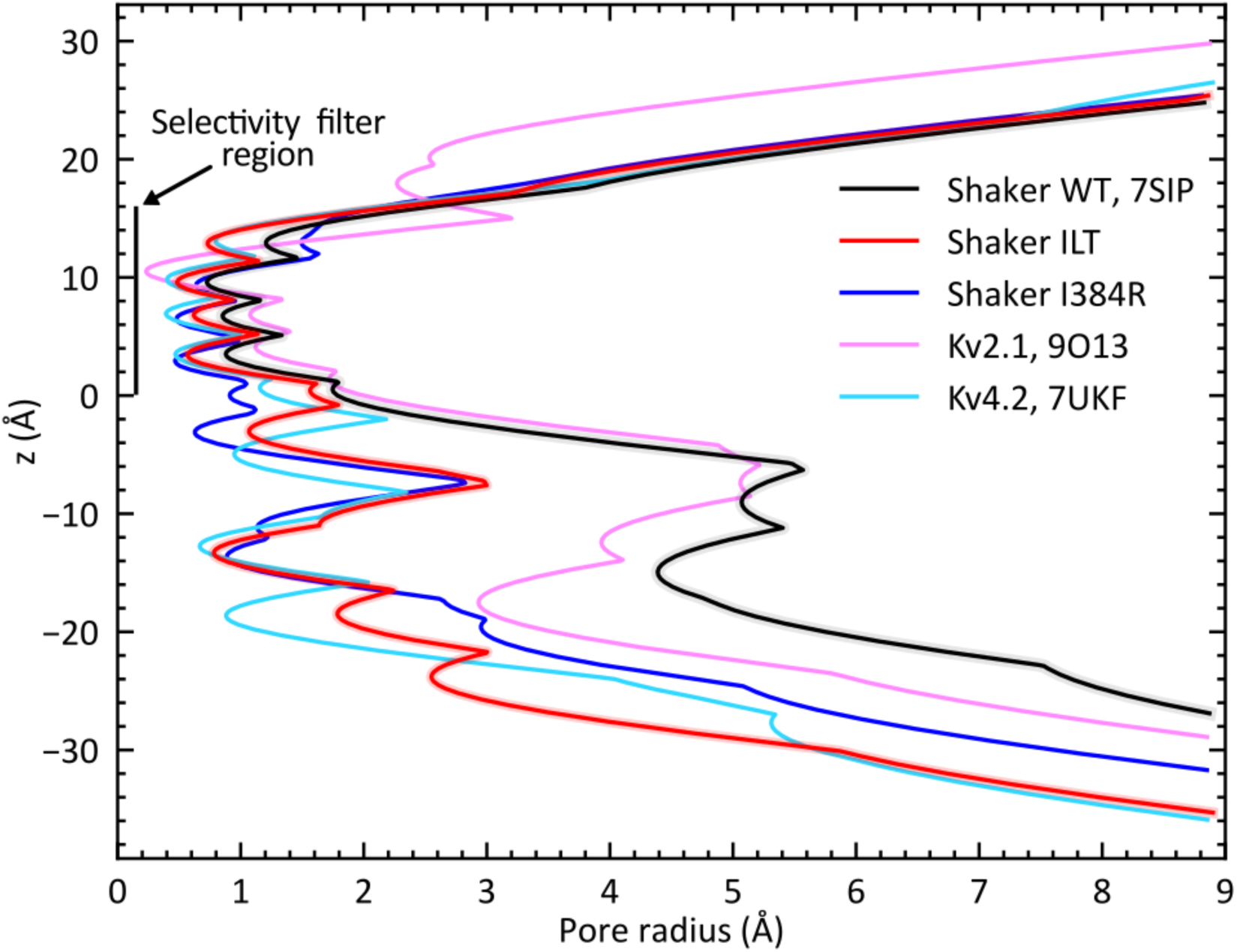
Comparison of pore radius. Shown are the pore radii of different Shaker/Kv structures from the Protein Data Bank (PDB). Those include Shaker WT (7SIP), Shaker ILT (9ZS7), Shaker I384R (9OIC), Kv2.1 (9O13), and Kv4.2 (7UKF).

**Figure S7.**
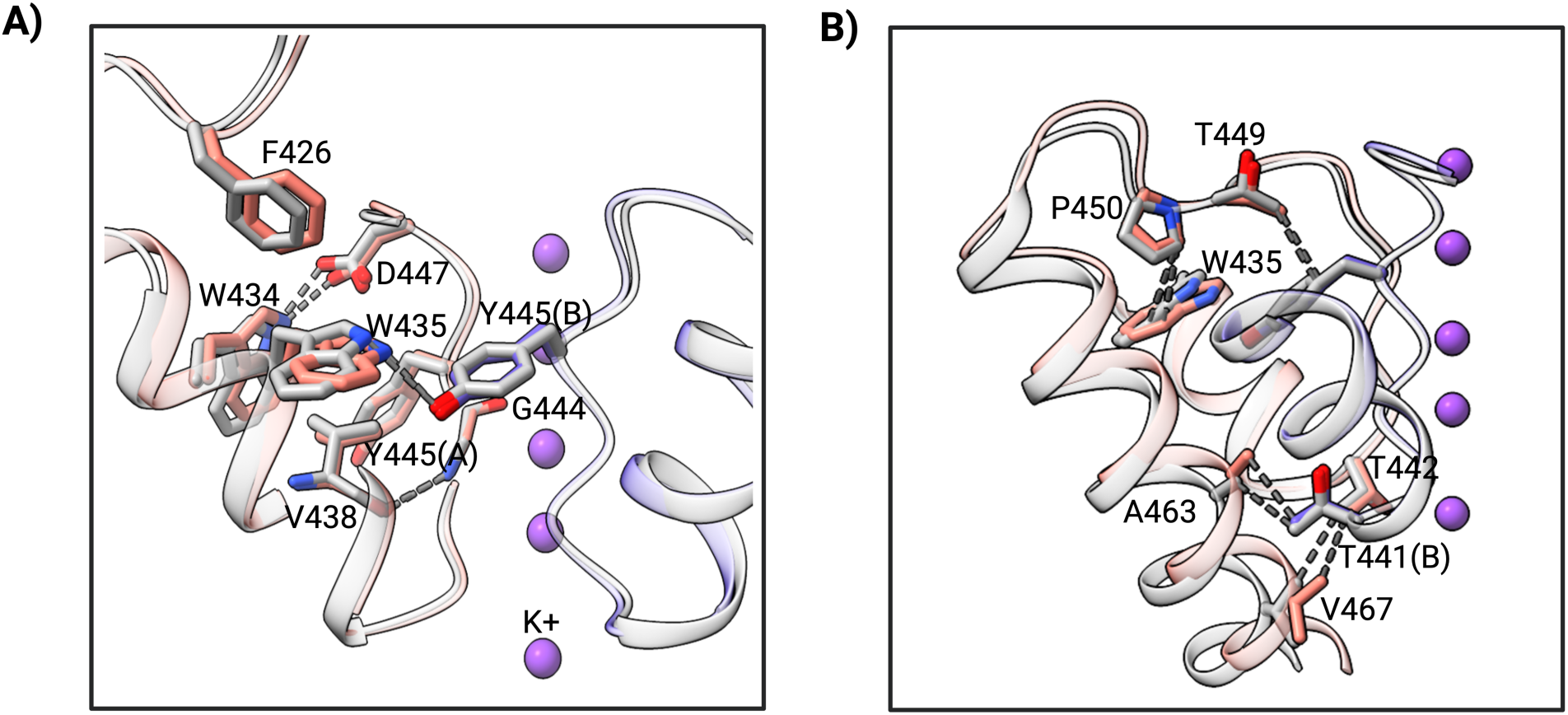
Selectivity filter interactions in Shaker ILT (colored) compared to WT (gray). **A)** Important hydrogen bond interactions (W434 (A chain) to D447 (A chain) and W435 (A chain) to Y445 (B chain)) are intact. **B)** A shift of VDW interactions occurs in T442 (chain A) to V467 (chain A) and T441 (chain B) to A463 (chain A).

**Figure S8.**
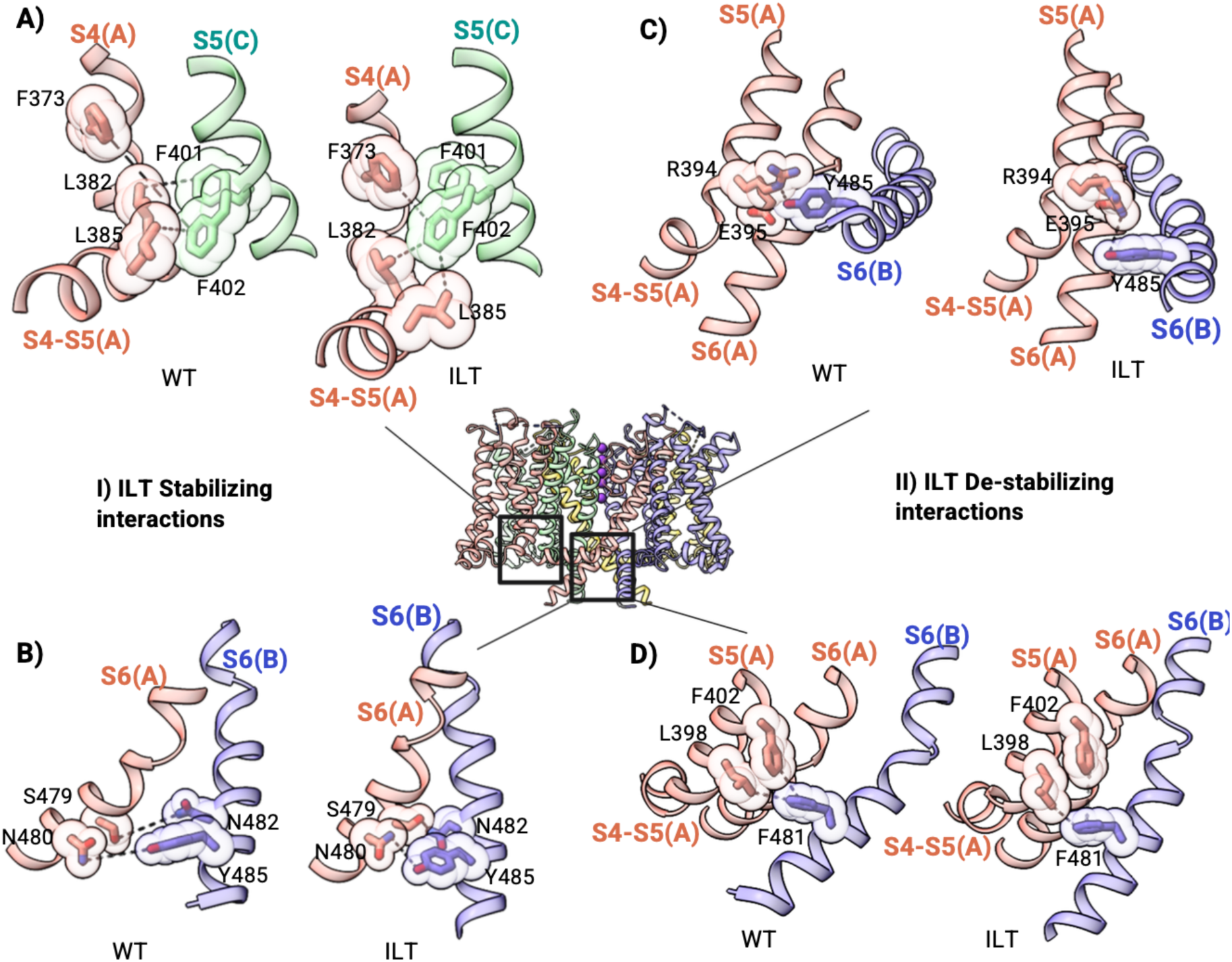
Interaction of intracellular gate. **A)** S4 helix interaction with S4-S5 linker in ILT compared to WT. Shift of S4 helix in ILT allows F373 stacking interactions with F401 and F402 in the S5 helix of chain C; L382 and L385 in the S4-S5 linker come closer to S5 helix of chain C in ILT compared to WT and form strong VDW interactions. **B)** S479 and N480 in S6 helix of chain A get closer in ILT compared to WT and form hydrogen bonds with N482 and Y485 in S6 helix of chain B. These interactions shown in **A** and **B** stabilize the ILT conformation. **C)** R394 and E395 in S4-S5 of chain A form a groove occupied by Y485 in S6 helix of chain B in WT. Y485 moves away from the groove in ILT. **D)** In WT, L398 and F402 of S5 helix of chain A form a groove occupied by F481 in S6 helix of chain B whereas in ILT F481 moves away from the groove. These interactions shown in **C** and **D** stabilize the WT conformation.

**Figure S9.**
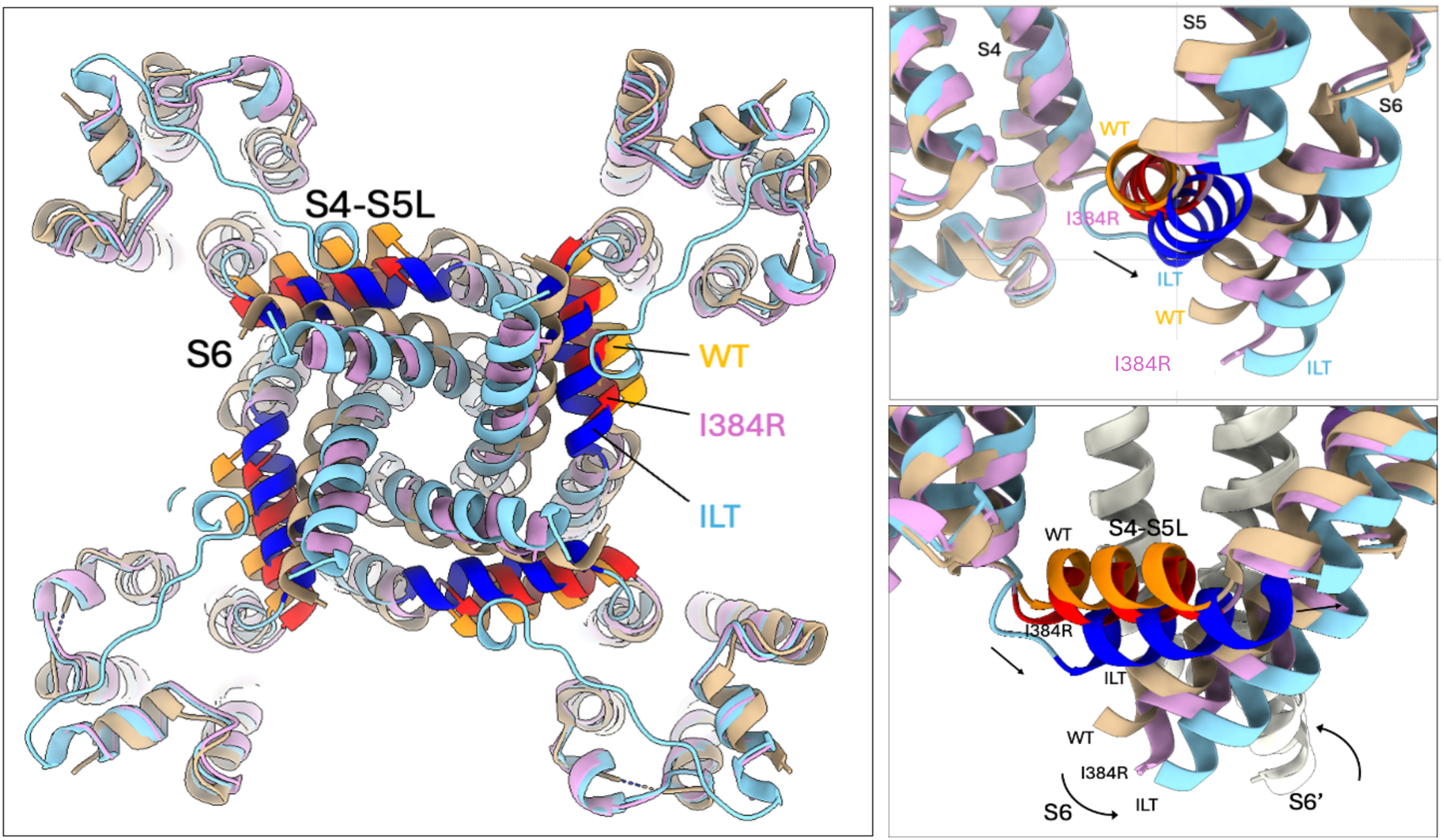
Comparison of the S4-S5 linker and S5-S6 gate in the WT, ILT and I384R experimental structures. Shaker ILT (PDB ID: 9ZS7) colored in cyan/blue, Shaker I384R (PDB ID: 9OIC) colored in pink/red (*37*), and WT (PDB ID: 7SIP) colored in gold/orange (*38*).

**Figure S10.**
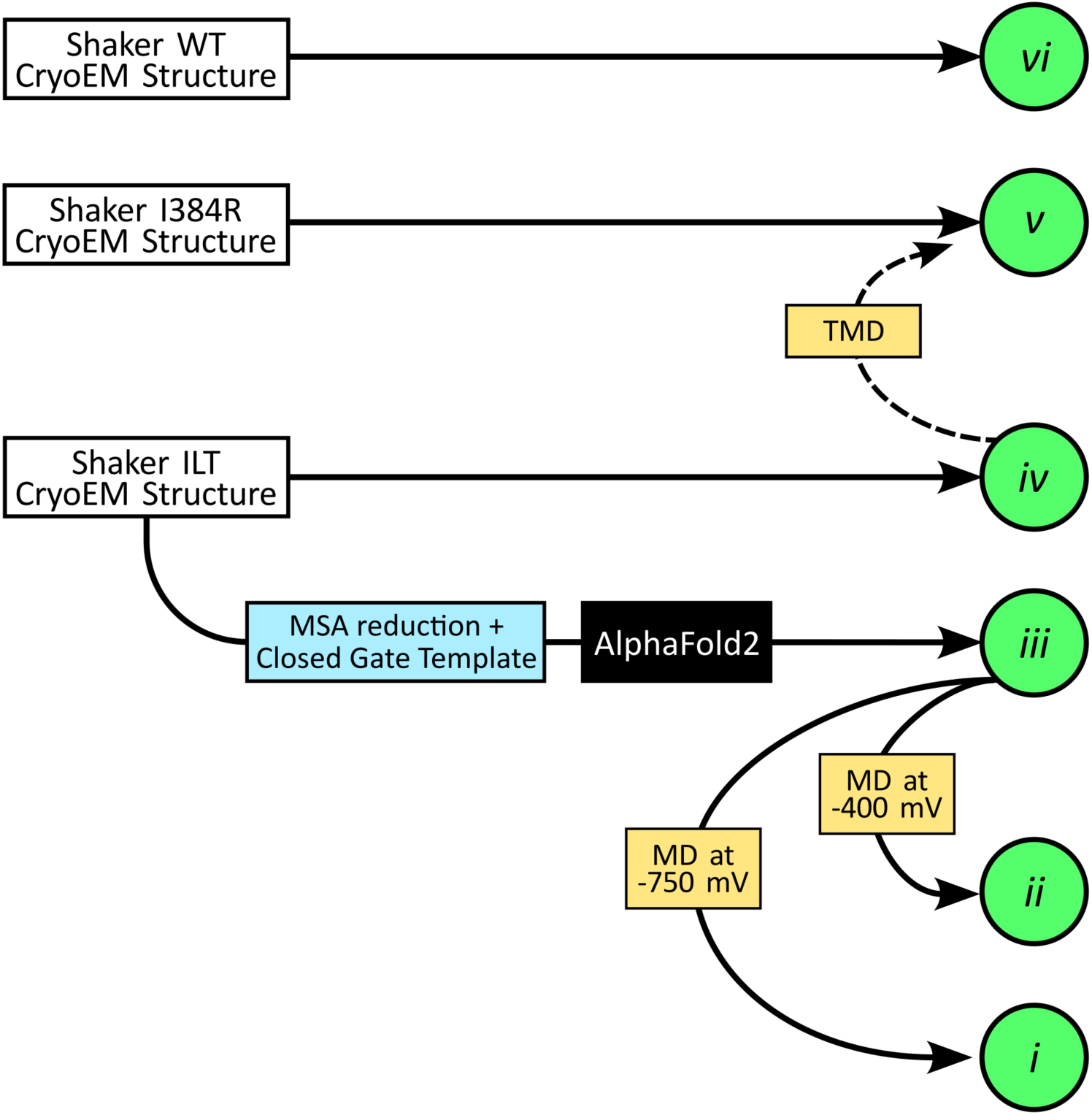
Structural modeling of missing states of the voltage activation process. The structural modeling relies on molecular dynamics (MD) simulations and predictions with AlphaFold2. The Shaker ILT cryo-EM structure was used as a template for AlphaFold2 and further MD was run on the AlphaFold predicted model. Hyperpolarizing voltages were used to obtain conformational states of the voltage sensor with an S4 helix progressively vertically shifted downwards. A model of state v was also obtained from a simulation of the structure of the Shaker mutant I384R (9OIC) after reverting to the WT sequence.

**Figure S11.**
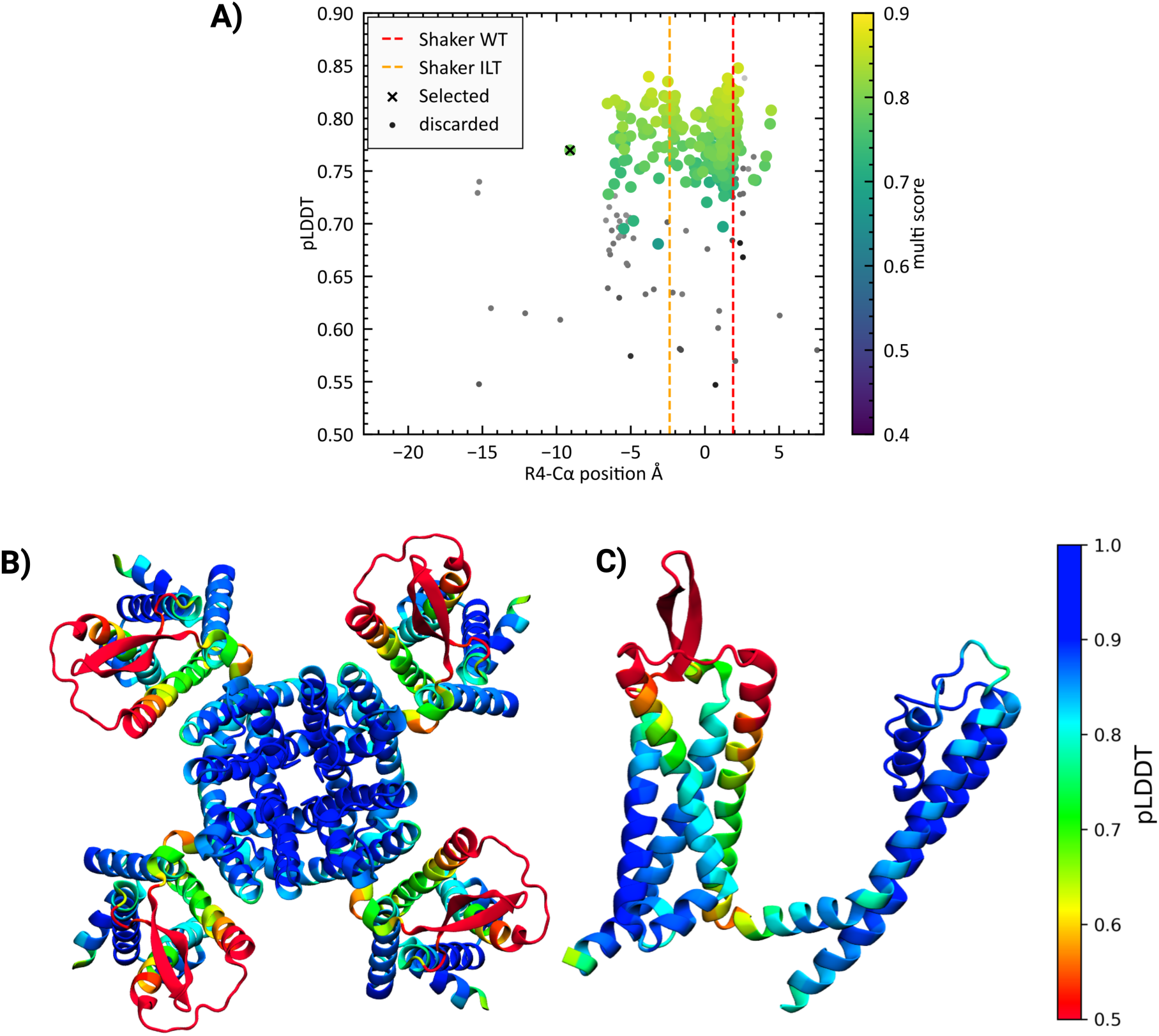
AlphaFold2 Score. **A)** Position of R4-Cα relative to Phe290-hydrophobic plug Cα vs Alphafold pLDDT score for Alphafold structures generated from both WT and ILT sequences. Colors indicate ‘multi’ score (0.8 × interchain pTM + 0.2 × pTM). Structures that were filtered out are shown in grayscale. Positions of the selected 2-click down model and experimental structures are shown. **B)** Per-residue pLDDT scores for the selected Alphafold 2-click down model, viewed as the tetramer (top view) and **C)** a single chain (front view).

**Figure S12.**
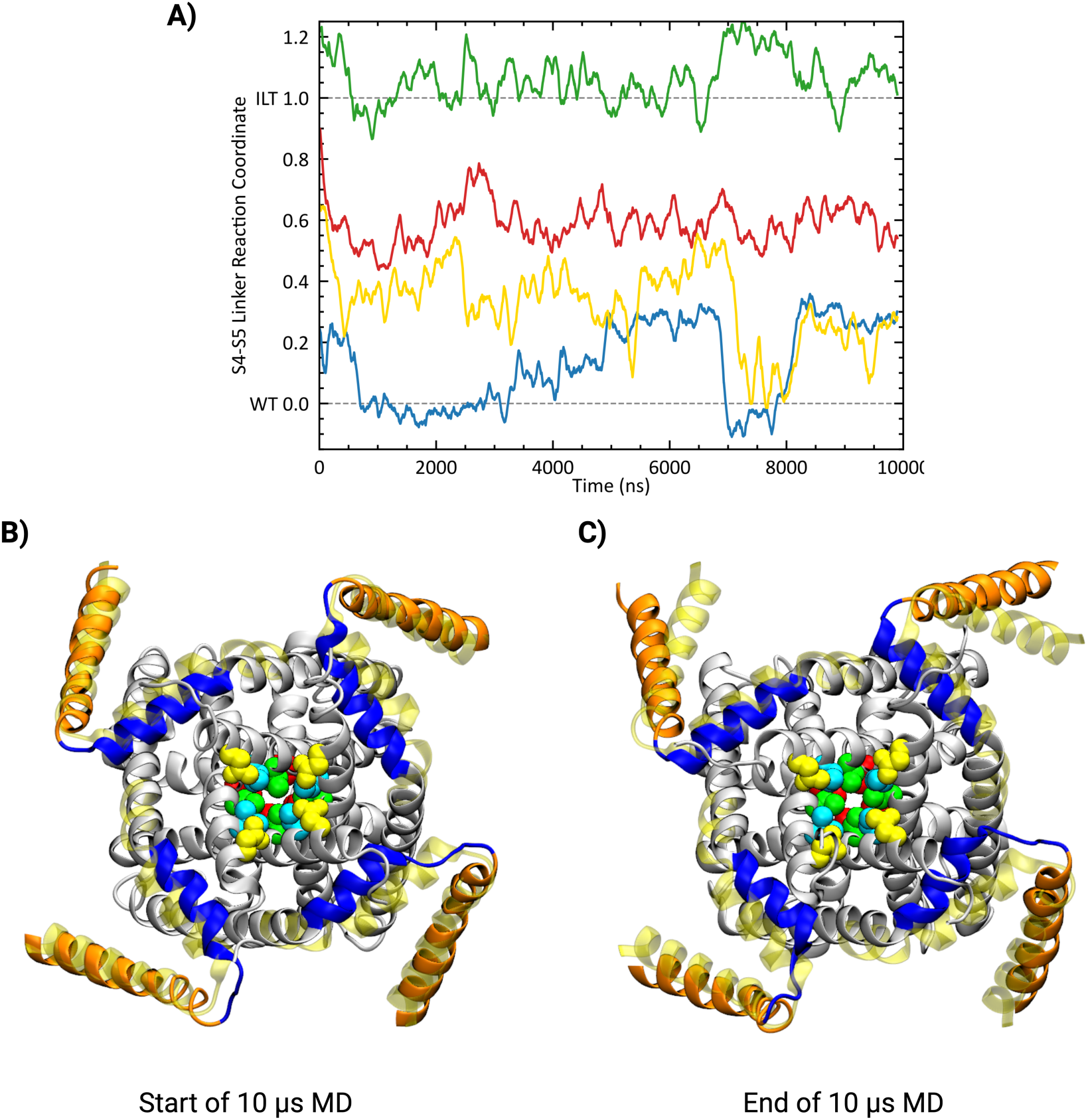
Dynamics of S4-S5 and intracellular gate in the state v (Up*-closed state). **A)** Behavior of S4-S5 helix of each subunit in 10 µs simulation of state v, shown as a projection onto a linear reaction coordinate of RMSD to WT (at 0) and ILT (at 1). Values shown are averaged over 100 ns. **B)** Snapshot at the start of the 10 µs trajectory **C)** Snapshot at the end of the 10 µs trajectory where the S4 helix is shown in orange and the S4-S5 linker is shown in blue. The hydrophobic residues at the intracellular gate (pore) which act as constriction points along the cavity are shown in red (Ile470), green (Val474), blue (Val478), and yellow (Asn482).

**Figure S13.**
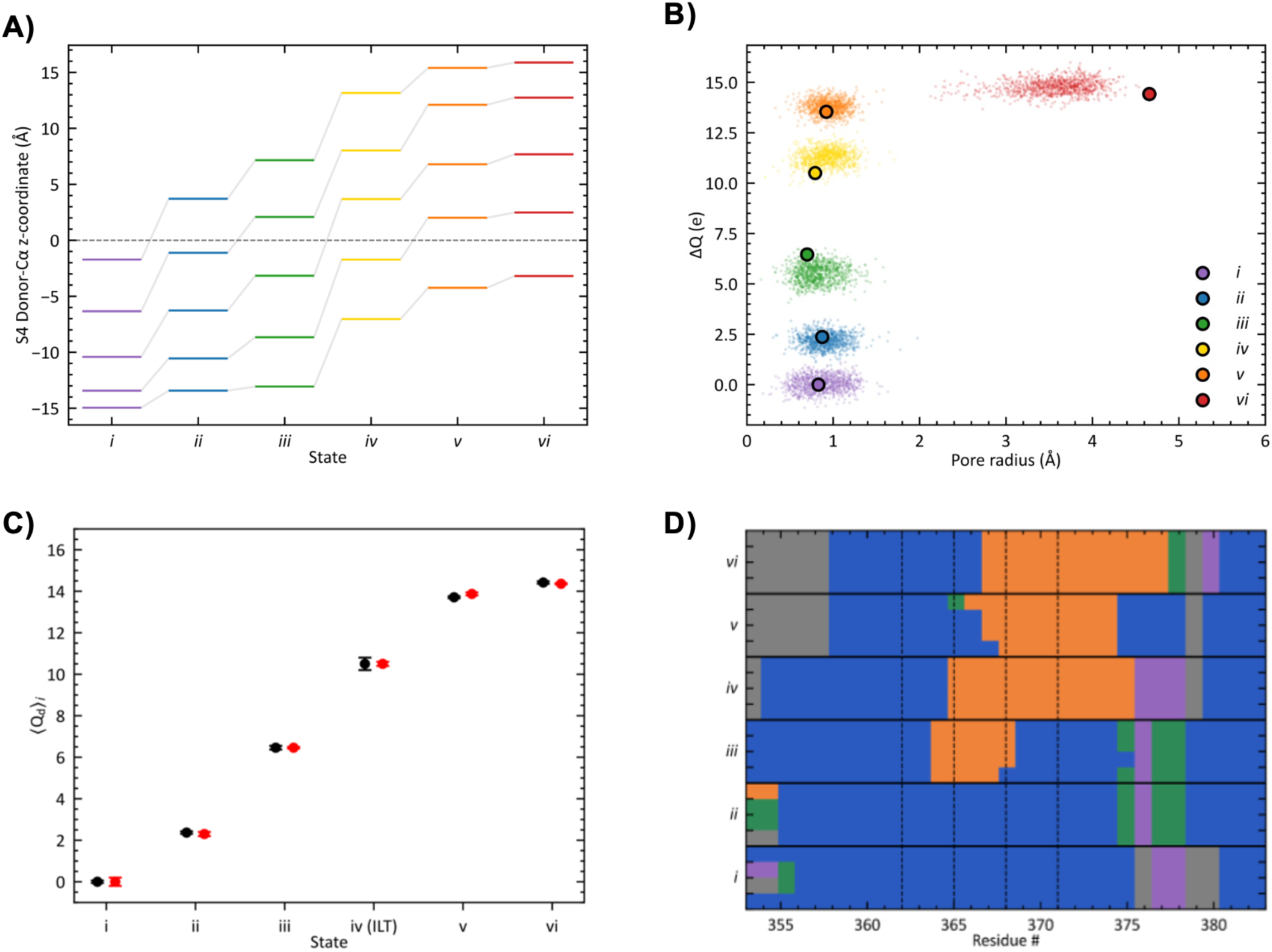
Voltage sensor vertical shift, gate opening, and secondary structure at S4 helix among the states. **A)** Vertical Cα shift of the gating charge Arg residues among the states along the activation process. **B)** Relating the opening of the inner gate by pore radius at the level of Val474 to the gating charge estimates of the states. The points in the background are the gating charge from the production MD simulations of each state. **C)** Sequence independence of the gating charge estimates between WT in black and ILT in red from the average displacement charge from MD simulations at zero membrane potential. **D)** Helicity of S4 helix in different structure models, where alpha helix is shown in blue, 3_10_-helix is shown in orange, hydrogen bonded turns in green, bends in purple, loops in grey.

**Figure S14.**
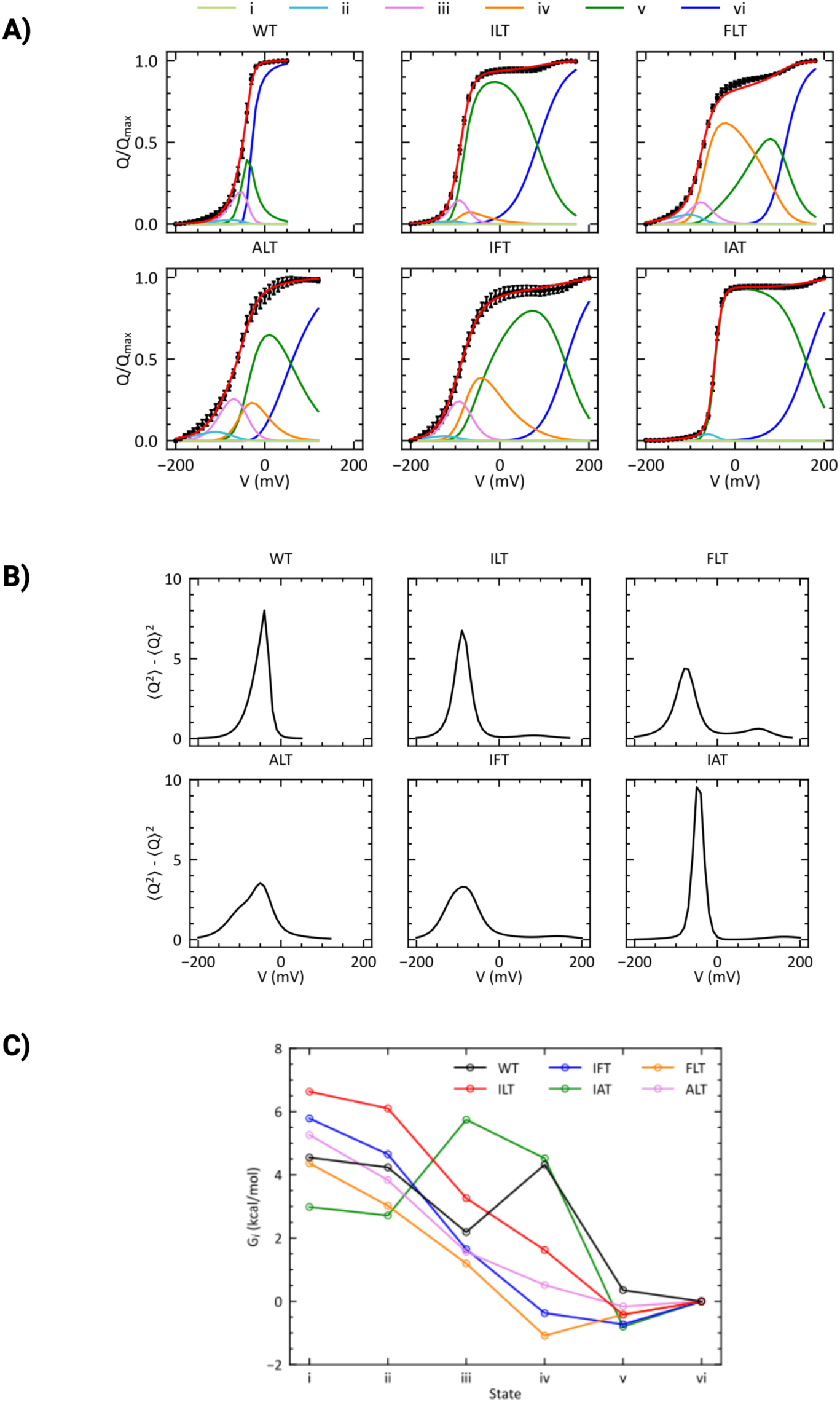
Fitting Q-V curves of Shaker mutants from Scheme 1. **A)** The fitting of the gating charge voltage dependence using Scheme 1. Scheme 1 shows the activation of Shaker which involves the transition through five closed states before opening. The first four transitions are modelled as subunit independent transitions, and the final transition is a cooperative opening of the inner gate. **B)** Voltage dependence of the variance of the gating charge from the model with the fitted parameters used for the Q-V. **C)** Relative free energy at zero membrane potential obtained from the Q-V curve fitting for the different mutants using the ΔQ of states *i* to *vi*.

**Figure S15.**
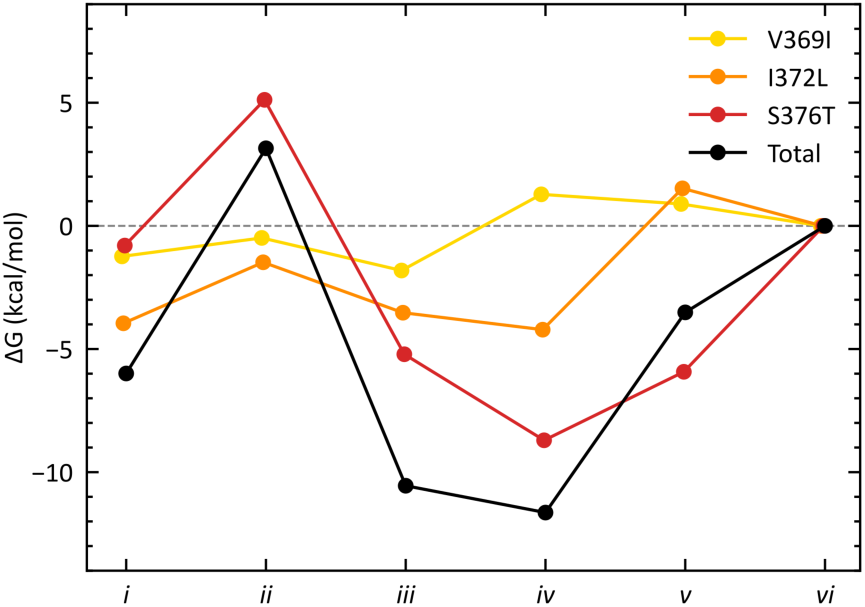
Average interaction of the ILT mutations and structural stability. Average change in interaction energy of ILT mutant relative to the WT calculated using MD simulations of each of the six conformational states. The last 10 ns of a 20 ns simulation was used for ILT and 50 ns was used for WT. The values are reported relative to the open active state. The average change in interaction energy is meant to approximate the ΔG from an alchemical free energy calculation. The ΔGs are also reported per residue and summed to a total for the triple mutant. Standard error of the total (not shown) is in the range of 5-12 kcal/mol.

**Table S1.**
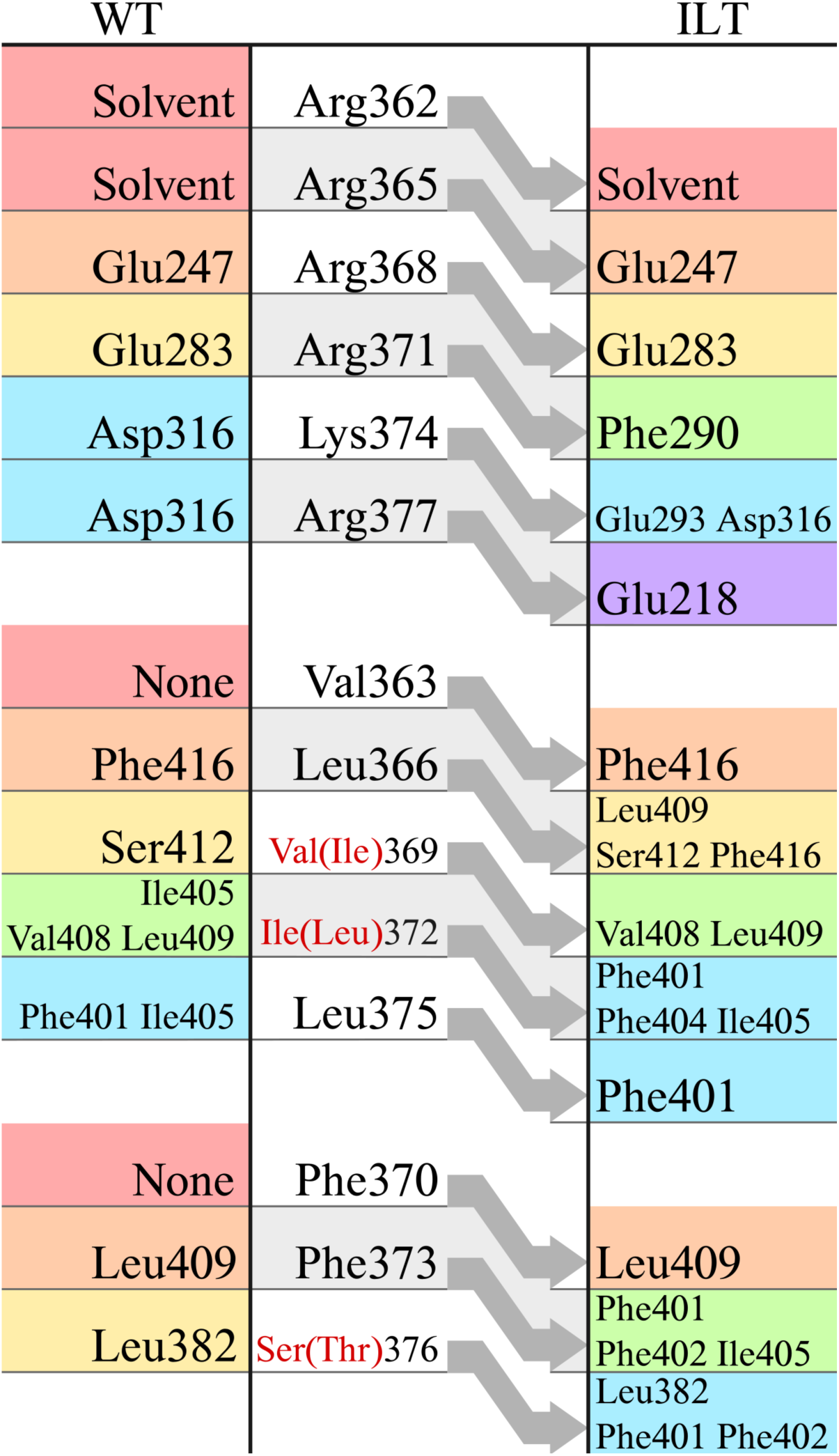
Interactions within the VSD.

